# Identifying Sequence Perturbations to an Intrinsically Disordered Protein that Determine Its Phase Separation Behavior

**DOI:** 10.1101/2020.01.06.894576

**Authors:** Benjamin S. Schuster, Gregory L. Dignon, Wai Shing Tang, Fleurie M. Kelley, Aishwarya Kanchi Ranganath, Craig N. Jahnke, Alison G. Simpkins, Roshan Mammen Regy, Daniel A. Hammer, Matthew C. Good, Jeetain Mittal

**Affiliations:** Department of Bioengineering, University of Pennsylvania, Philadelphia, PA, 19104; Department of Chemical and Biochemical Engineering, Rutgers University, Piscataway, NJ 08854; Department of Chemical and Biomolecular Engineering, Lehigh University, Bethlehem, PA 18015; Laufer Center for Physical and Quantitative Biology, Stony Brook University, Stony Brook, NY, 11794; Department of Physics, Brown University, Providence, RI, 02912; Department of Chemical and Biomolecular Engineering, University of Pennsylvania, Philadelphia, PA, 19104; Department of Cell and Developmental Biology, University of Pennsylvania, Philadelphia, PA, 19104

## Abstract

Phase separation of intrinsically disordered proteins (IDPs) commonly underlies the formation of membraneless organelles, which compartmentalize molecules intracellularly in the absence of a lipid membrane. Identifying the protein sequence features responsible for IDP phase separation is critical for understanding physiological roles and pathological consequences of biomolecular condensation, as well as for harnessing phase separation for applications in bio-inspired materials design. To expand our knowledge of sequence determinants of IDP phase separation, we characterized variants of the intrinsically disordered RGG domain from LAF-1, a model protein involved in phase separation and a key component of P granules. Based on a predictive coarse-grained IDP model, we identified a region of the RGG domain that has high contact probability and is highly conserved between species; deletion of this region significantly disrupts phase separation in vitro and in vivo. We determined the effects of charge patterning on phase behavior through sequence shuffling. By altering the wild-type sequence, which contains well-mixed charged residues, to increase charge segregation, we designed sequences with significantly increased phase separation propensity. This result indicates the natural sequence is under negative selection to moderate this mode of interaction. We measured the contributions of tyrosine and arginine residues to phase separation experimentally through mutagenesis studies and computationally through direct interrogation of different modes of interaction using all-atom simulations. Finally, we show that in spite of these sequence perturbations, the RGG-derived condensates remain liquid-like. Together, these studies advance a predictive framework and identify key biophysical principles of sequence features important to phase separation.

**Significance Statement:** Membraneless organelles are assemblies of highly concentrated biomolecules that form through a liquid-liquid phase separation process. These assemblies are often enriched in intrinsically disordered proteins, which play an important role in driving phase separation. Understanding the sequence-to-phase behavior relationship of these disordered proteins is important for understanding the biochemistry of membraneless organelles, as well as for designing synthetic organelles and biomaterials. In this work, we explore a model protein, the disordered N-terminal domain of LAF-1, and highlight how three key features of the sequence control the protein’s propensity to phase separate. Combining predictive simulations with experiments, we find that phase behavior of this model IDP is dictated by the presence of a short conserved domain, charge patterning, and arginine-tyrosine interactions.

## Introduction

Liquid-liquid phase separation (LLPS) of biomolecules is a highly robust and ubiquitous phenomenon in biology, enabling compartmentalization in the absence of delimiting membranes^1^. Biomolecular LLPS commonly occurs within the cell, forming compartments that have been termed biomolecular condensates or membraneless organelles^2^ and include stress granules^3–5^, P-granules^1, 6^, nucleoli^7^, and numerous others^8–13^. Most membraneless organelles contain an overrepresentation of proteins with intrinsically disordered and low-complexity regions^14^, which are important drivers of phase separation behavior^15, 16^. Therefore, decoding the sequence determinants of intrinsically disordered protein (IDP) phase separation is important for understanding the biochemistry of biomolecular condensates in physiological and pathophysiological conditions. Characterizing the effects of sequence on phase behavior is also important for the field of protein-based materials^17^, wherein proteins can be designed to have desired characteristics and programable assembly^18–20^, with applications in biotechnology such as drug delivery, cell engineering, and biomimetics^21–24^.

Here we investigate a model IDP sequence from LAF-1, which is a member of the DDX3 family of RNA helicases and is a major component of P-granules, membraneless organelles involved in germline specification in *C. elegans* embryos^25^. LAF-1 contains an N-terminal domain of 168 residues that is intrinsically disordered, followed by a folded helicase domain, and a short disordered prion-like domain at the C-terminus^6^. The N-terminal domain contains an abundance of glycine and arginine residues, with several occurrences of the motif RGG, and is hereafter referred to as LAF-1 RGG. Importantly, the RGG domain is necessary and sufficient for phase separation^6^, although both experimental and computational studies have shown that inclusion of the folded domain increases the protein’s ability to phase separate^26, 27^.

LAF-1 RGG is an excellent model system for exploring the sequence determinants of protein phase separation because it is believed to be fully disordered, and it contains a sufficient diversity of amino acids to enable different types of interactions^28, 29^. The advantage of a fully disordered sequence is that it allows for relatively distributed interactions between all residues, so the relationship between amino acid composition and phase behavior can be more readily ascertained, as compared to proteins with residues buried in folded domains. LAF-1 was one of the first proteins found in biomolecular condensates in vivo and whose phase behavior was mapped in vitro, yet key questions remain about its properties and function ^6, 27^. Additionally, we have recently designed constructs based on LAF-1 RGG to generate micrometer-sized protein condensates that can respond to specific stimuli and that can selectively compartmentalize cargo proteins, progressing toward the design of synthetic organelles that may be expressed in cells and that are orthogonal to normal cellular function ^21^. To advance the design of synthetic organelles, we seek to understand how perturbations to the RGG domain sequence may alter phase behavior in a predictableway^18, 30^.

In this work, we use simulations and experiments to characterize the sequence-dependent LLPS of the LAF-1 RGG domain, identifying perturbations that result in significant changes to the phase behavior. First, we have identified a small hydrophobic region that exhibits high contact probability in coarse-grained (CG) molecular dynamics simulations, and that contains a well-conserved specific binding site for the eukaryotic translation initiation factor 4E (eIF4E)^31^. We demonstrate that removal of this region greatly reduces the phase separation propensity of the RGG domain in silico, in vitro, and in vivo in a eukaryotic model, suggesting that the hydrophobic interactions within this region are also important to LLPS. Second, we show that shuffling the amino acid residues of the RGG sequence to introduce charge patterning can drastically increase phase separation propensity and that by simultaneously preserving the conserved hydrophobic region, we can further increase it. Third, we investigate alterations to amino acid composition by mutating tyrosine to phenylalanine and arginine to lysine, mutations that affect LLPS propensity of FUS^30, 32^. We find that tyrosine to phenylalanine and arginine to lysine mutations both reduce the phase separation propensity of the LAF-1 RGG domain. We then identify the interaction mechanisms disrupted by these mutations as being hydrogen bonds, cation-π interactions, and sp^2^/π interactions, all three of which are present between arginine and tyrosine and may act cooperatively, whereas at least one of these is impossible upon mutation. Importantly, we rule out a previous model based exclusively on arginine-tyrosine interaction, which cannot predict the critical concentration for LAF-1 RGG phase separation. Finally, we show that the RGG-derived condensates remain liquid-like despite these three classes of sequence perturbations, indicating that phase behavior can be tuned independent from material properties. Our combined results elucidate fundamentally new and important sequence determinants of IDP phase separation while demonstrating a computationally-guided approach for studying phase behavior of biomolecular condensates. These results promise a framework toward the rational design of LLPS-enabled IDPs.

## Results

### A short, conserved, hydrophobic region is important for LLPS of the RGG domain

We focused our efforts on the RGG domain of LAF-1, as it is necessary and sufficient to drive phase separation^6^, making it an ideal model system to understand the sequence determinants of LLPS. Phase separation of LAF-1 RGG is hypothesized to be driven by several different modes of interaction, including electrostatic, π-π, and cation-π interactions^27^. In addition, hydrogen bonds and hydrophobic contacts may play a role in phase separation for sequences containing residues capable of such interactions^16, 33–37^. However, it is difficult to characterize these interactions using experimental techniques due to the dynamic nature of the phase-separated proteins and the high spatiotemporal resolution needed to probe the interactions^35^.

To provide insight into the sequence determinants of phase separation, we conducted simulations of a condensed assembly of 100 chains of LAF-1 RGG using a transferrable CG model (see Methods), which accounts for the combined interaction modes between each amino acid pair^26^. The condensed assembly is liquid-like, with chains exhibiting liquid-like diffusion, as we have shown in the previous work^26^. We then enumerated the average number of intermolecular contacts formed between each residue of the sequence with each residue in all other protein chains, which may represent many different modes of interaction at the atomic scale. The results highlighted a single region spanning residues 21-30 (RYVPPHLRGG) having highly enhanced contact probability within the condensed protein assembly (Fig. 1A). This region has a considerably different composition from the full RGG sequence, particularly since it contains several hydrophobic residues: this region contains the only two Pro, the only Val, and one of the only two Leu in the entire RGG domain. Region 21-30 is more prone to interaction, not only with itself but also with many regions of the protein (Fig. 1A).

**Figure 1:**
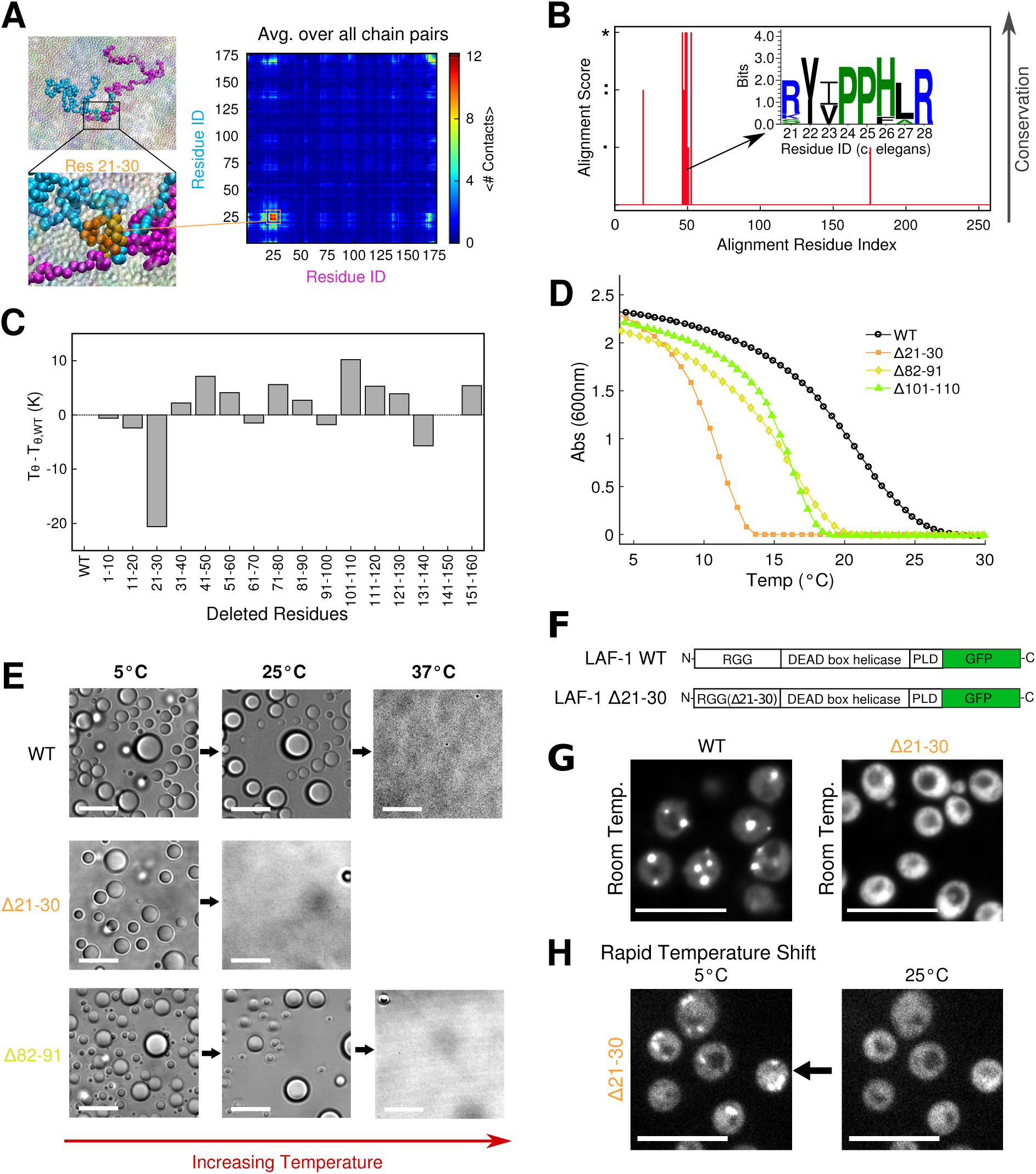
A short segment of LAF-1 RGG is critical for phase separation: A) Coarse-grained sequence-specific simulations of LAF-1 RGG highlight a small region where contact probability is enhanced. Insets show the interaction of two protein chains and zoomed view of contacts between residues within the contact-prone region. B) Sequence analysis of LAF-1 and some of its homologs highlight high sequence conservation in the folded helicase domain, and poor conservation in the disordered RGG and prion-like domains (Fig. S1A). Within the RGG domain, we identify one short region having good conservation, which corresponds to the region highlighted by CG simulations. The amino acids within the sequence are displayed as an inset logo. C) Results of deleting 10 amino segments, scanning across the sequence of RGG; Tθ from CG simulations. Errors are very small and would not show up well on the bar plot. D) Turbidity measurements show temperature-dependent phase behavior of WT RGG vs. variants with deletion of residues 21-30, 82-91, or 101-110. Proteins phase separate upon cooling from above to below the phase transition temperature. Protein concentrations were 1 mg/mL (approximately 60 μM) in 150 mM NaCl buffer, pH 7.5. Data shown is representative of three independent turbidity experiments for each protein (Fig. S2). Similar to previous work^18^, we have not averaged the repeats, and therefore, we have not added error bars because the temperatures of the measurements from different replicates are not exactly the same. E) RGG Δ21-30 and RGG Δ82-19 condense into spherical liquid droplets, similarly to WT RGG, as shown by brightfield microscopy. Upon heating from 5 °C, RGG Δ21-30 droplets dissolve at a lower temperature compared to WT or RGG Δ82-91. Protein concentration and buffer are the same as for turbidity assay. Scale bars: 10 μm. F) Schematic for full-length LAF-1 constructs including C-terminal GFP fluorescent tag. (PLD: prion-like domain.) G) Full-length LAF-1 phase separates in yeast at room temperature, with multiple puncta per cell. In contrast, LAF-1 Δ21-30 does not phase separate at room temperature; delocalized fluorescence in the cytoplasm is observed. H) Upon sufficient cooling, LAF-1 Δ21-30 does exhibit phase separation in yeast: fluorescent condensates form rapidly upon cooling from 25 °C to 5 °C, consistent with in vitro results in (D). Scale bars: 10 μm.

Interestingly, subregion 21-28 corresponds exactly with the previously identified eIF4E-binding motif^38^. We conducted a homology search, which also confirmed this region as an important functional motif due to its high degree of conservation across diverse species (Fig. 1B, S1A; Supporting Methods). The level of conservation is likely due to its biological function, rather than its importance to phase separation per se. However, the presence of a domain prone to self-association will still make considerable contributions to phase separation^39^. We were curious whether this region alone would undergo LLPS, and thus conducted CG simulations on just the 8-residue fragment. Due to the small chain length and net charge, we were unable to observe LLPS for the fragment alone, even at very high concentrations and low temperatures (Movie S1).

We, therefore, conducted simulations to compare how deletion of residues 21-30 vs. other regions of the RGG domain affect phase behavior to gain additional insight into the extent to which different regions of the RGG domain contribute to phase separation. Previously, we have shown that the θ-temperature (Tθ), where a single IDP chain behaves as in an ideal solvent, can serve as a good proxy for the critical temperature of phase separation (Tc)^40^, above which the IDP will always form a single continuous phase regardless of the protein concentration. Taking advantage of this relationship, we tested the effects of deleting distinct 10-residue segments from the LAF-1 RGG sequence by conducting single-chain simulations across a range of temperatures. We identified Tθ for each deletion and how it deviates from that of the WT RGG sequence (Fig. 1C). The Δ21-30 variant shows the greatest reduction of θ-temperature, indicating that it would have the lowest propensity to phase separate. In contrast, most other deletions had little effect or actually raised Tθ. This strongly suggests that the sticky hydrophobic subregion has an important role in phase separation of the LAF-1 RGG domain. We note that many of the deletion sequences have a higher Tθ than the full-length RGG, counter to the expectation that longer chain length generally favors LLPS. We believe this effect in the simulation model can be attributed to a subtle balance between the changes in hydrophobicity, net charge, and SCD rather than a single sequence descriptor (Figure S1B). Given the simplicity of our simulation model and the errors associated with predicting phase separation based solely on Tθ, it is possible our computational framework can distinguish sequences such as Δ21-30 which have more significant changes to LLPS behavior, but cannot capture smaller changes as with the other sequences.

We then tested these predictions experimentally by recombinantly expressing and purifying RGG and its variants (Fig. S1C, S1D). To study protein phase behavior, we used a temperature-dependent turbidity assay, in which protein solutions are cooled from above to below their phase transition temperature. Proteins transition from well-mixed to demixed upon cooling below the saturation temperature (T_sat_), defined as the point where we first observe an increase in the measured solution turbidity from that of the well-mixed solution. WT RGG and the deletion variants all exhibited upper-critical solution temperature phase behavior, becoming turbid upon cooling (Fig. 1D, S2A), characteristic of IDPs rich in polar and charged amino acids^18, 41^. Under these experimental conditions, the T_sat_ of WT RGG is approximately 26 °C, whereas the variant with the sticky hydrophobic subregion deleted (Δ21-30) has a phase transition temperature of only approximately 14 °C, representing a decrease of 12 °C. We tested two additional deletion variants, the first having residues 101-110 deleted (Δ101-110) which displayed the highest Tθ value according to simulations (Fig. 1C), and a control sequence having residues 82-91 deleted (Δ82-91), which contains the same number of arginine and tyrosine residues as Δ21-30. Both of these display a more modest reduction of T_sat_, by roughly 6 °C. These results indicate that the eIF4E-binding motif has the effect of promoting phase separation of the LAF-1 RGG domain, in addition to its specific binding function.

We then assessed whether the turbidity was due to the formation of spherical droplets, a hallmark of LLPS. We employed an optical microscope equipped with a temperature controller capable of rapidly setting the sample temperature to above or below room temperature. Indeed, we observed that both WT and the deletion variants of RGG assembled into spherical droplets below their respective values of T_sat_. At low temperature (5 °C), Δ21-30, and the control deletions formed micrometer-scale liquid droplets that were morphologically indistinguishable from those formed by WT RGG (Fig. 1E). Notably, Δ21-30 droplets dissolved within 1 minute upon increasing the sample temperature from 5 °C to 25 °C, whereas Δ82-91 and WT RGG exhibited slower and incomplete droplet dissolution at 25 °C, requiring a temperature of 37°C to rapidly and fully dissolve (Fig. 1E). In all cases, the process was reversible in that droplets were able to assemble, disassemble, and reassemble upon cycling the temperature (Fig. S3). Thus, both the macroscopic turbidity assays and microscopy confirmed that purified Δ21-30 phase separates, but with significantly reduced phase separation propensity as compared to WT RGG and the other deletion variants. Finally, we assessed the effect of these deletions on the phase behavior of LAF-1 in living cells.

For these experiments, we selected *S. cerevisiae*, a well-established model for studying protein aggregation^42, 43^, and we used full-length GFP-tagged LAF-1 (Fig. 1F). At room temperature, we observed multiple bright cytoplasmic puncta in cells expressing WT LAF-1, whereas we observed only delocalized cytoplasmic fluorescence for LAF-1 Δ21-30 (Fig. 1G). We confirmed by western blot that WT LAF-1 and LAF-1 Δ21-30 expressed at similar levels (Fig. S1E). The full-length Δ21-30 variant rapidly formed fluorescent cytoplasmic puncta when cooled to 5 °C, which then rapidly dispersed at 25 °C (Fig. 1H). This suggests that residues 21-30 are indeed important for phase separation of full-length LAF-1 in living cells, with their deletion resulting in LAF-1 having a reduced propensity to phase separate. While deletion of this region would also likely impact the interactions of LAF-1 with eIF4E, the appreciable difference observed in the simulations and *in vitro* experiments – which do not incorporate the eIF4E protein – indicate that the eIF4E-binding motif itself is contributing to phase separation. It will be interesting to consider in the future how the position of the eIF4E binding region within the disordered LAF-1 RGG domain, in the context of the full-length protein, may affect its phase behavior and function.

In total, our in vitro and in vivo results suggest that LAF-1 phase separation is driven by multivalent interactions in addition to strong interactions with the more hydrophobic eIF4E-binding motif. Although this 10 amino acid motif is necessary, it is not sufficient to control RGG phase separation, and therefore we sought additional sequence determinants.

### Charge distribution and sequence shuffling can be used to control LLPS

We next sought to understand how the patterning of amino acids can influence the phase separation of LAF-1 RGG, as has been studied previously for other proteins^44, 45^, and the joint contributions of charge-charge interactions and the sticky hydrophobic subregion. We constructed one set of sequences having identical amino acid composition to WT RGG, but with the full sequence randomly shuffled, and a second set in which the eIF4E-binding motif (residues 21-28) was preserved. To quantify the extent to which we can expect the sequences to differ, we calculated the sequence charge decoration (SCD) parameter, where a more negative SCD score indicates greater charge segregation for sequences with many positive and negative charges. SCD has been shown to be correlated with disordered proteins’ radii of gyration (Rg)^46^, and with their critical temperatures (Tc)^44^.

To observe the accessible SCD space of polypeptides having the same composition as the LAF-1 RGG domain, we generated 1 million randomly shuffled sequences of LAF-1 RGG and plotted the probability distribution of SCD (Fig. 2A). We find that randomly shuffled sequences tend to populate a very small window of SCD values, with 93.6% of the shuffled sequences having SCD scores between -2 and 0.5. For comparison, the minimum possible value for a sequence of the same length and composition is -28.03, when following the constraints set by experimental procedures (Supporting Methods). Notably, the WT RGG sequence does not sit at the center of the distribution, but rather, its SCD (0.565) is in the highest 2% of the million randomly generated sequences. This is in contrast to the IDRs of similar helicase proteins such as DDX4, which is more charge-segregated^47^, having an SCD value of -1.02. Charge patterning could perhaps regulate phase separation *in vivo* such that the saturation concentration of LAF-1 is similar to that of the native expression level, and also make it distinguishable from other proteins of similar amino acid composition^48^.

**Figure 2:**
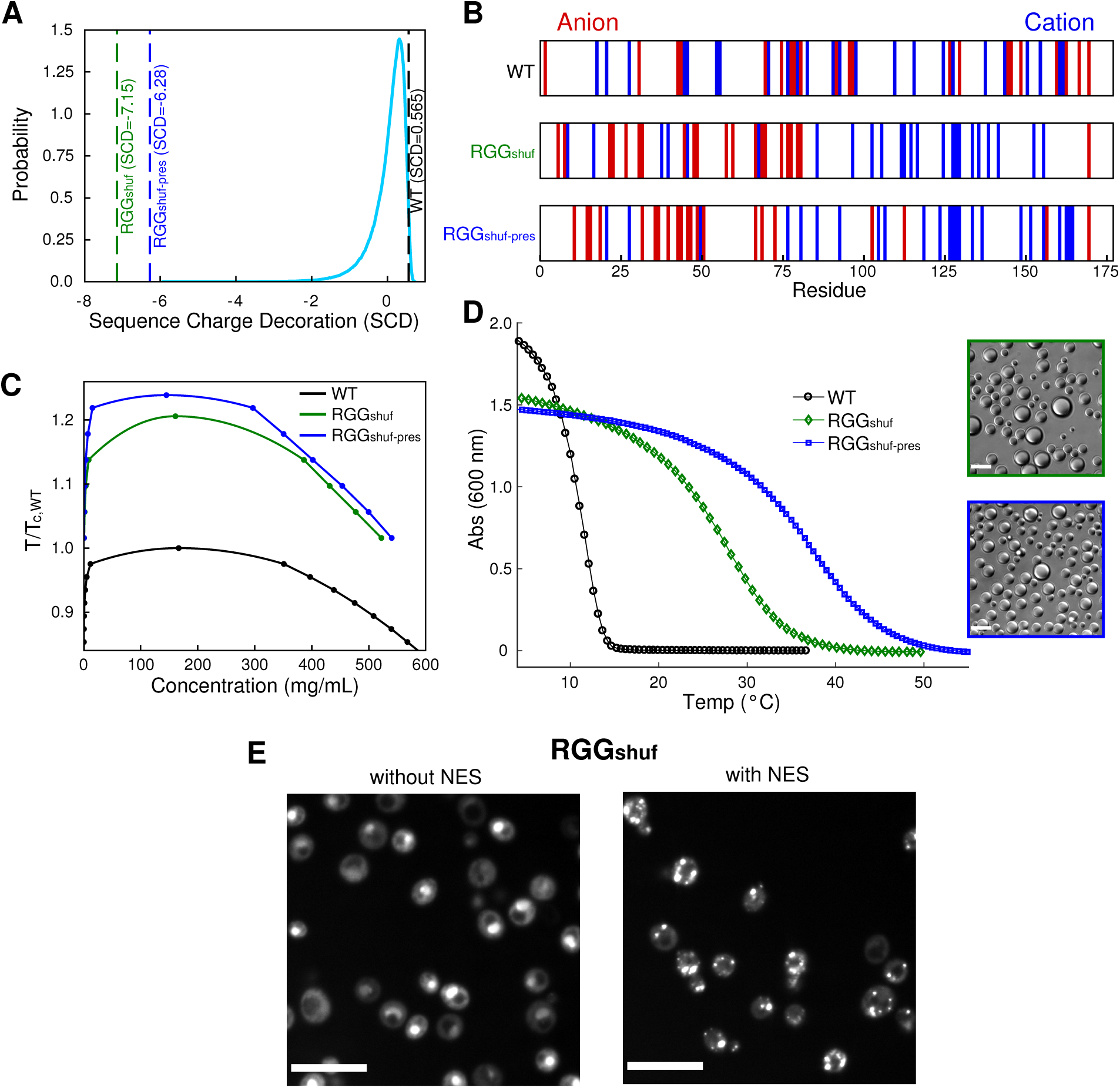
Charge patterning alters LAF-1 RGG phase transition: A) Probability distribution of sequence charge decoration (SCD) values from 1 million random shuffles of LAF-1 RGG. SCD values of WT, RGG_shuf_, and RGG_shuf-pres_ are highlighted with dashed lines. B) Location of charged residues in the three sequences. C) Phase diagrams of WT, RGG_shuf_, and RGG_shuf-pres_ from CG simulations. Temperatures are normalized to the critical temperature of WT RGG. Errors on the concentration axis are smaller than symbols. D) Turbidity measurements show the temperature-dependent phase behavior of WT RGG vs. RGG_shuf_ and RGG_shuf-pres_ variants. Data shown are representative of three independent turbidity experiments for each protein (Fig. S2). Protein concentrations were 0.3 mg/mL (approximately 17 μM) in 150 mM NaCl buffer, pH 7.5. Both RGG_shuf_ and RGG_shuf-pres_ exhibited phase transition temperatures markedly higher than that of WT RGG, and both appeared as liquid droplet condensates under optical microscopy at room temperature (insets; scale bars are 10 μm). E) LAF-1shuf-GFP expression in yeast. Charge patterning leads to constitutive import. The addition of NES enables LAF-1shuf-GFP to be cytosolic, and this variant exhibits protein condensate formation. Scale bars: 10 μm.

We selected the sequence with the lowest SCD value, termed RGG_shuf_. We did the same for sequences having the eIF4E-binding motif preserved (RGG_shuf-pres_), to test whether there is an appreciable difference between charge-segregated variants with and without the presence of a sticky hydrophobic subregion. The two sequences are depicted in Fig. 2B, which shows that both have an abundance of anionic residues in the first half of the sequence, and an abundance of cationic residues in the second half, in contrast with the WT sequence, which has a relatively even distribution of cationic and anionic residues throughout. We conducted CG molecular simulations for these sequences and determined the phase diagrams as a function of temperature. Both shuffled sequences show a drastic increase in the critical temperature compared to WT (Fig. 2C), as well as compaction in single-chain simulations (Fig. S4). Interestingly, RGG_shuf_ does not exhibit as large of an upward shift in Tc as does RGG_shuf-pres_, even though it has a slightly lower SCD value. This indicates that charge patterning is capable of inducing large shifts to the phase diagram, but a combination of charge segregation and preservation of the hydrophobic subregion promotes LLPS even more.

We then tested these predictions experimentally by conducting temperature-dependent turbidity assays on recombinantly expressed and purified WT RGG, RGG_shuf_, and RGG_shuf-pres_. These experiments were performed using lower concentrations (0.3 mg/mL) of protein because the two shuffled variants display a much greater propensity to phase separate (Fig. 2D, S2B). Remarkably, whereas WT RGG undergoes LLPS at approximately 15°C under these conditions, RGG_shuf_ demixed at 42°C, and RGG_shuf-pres_ demixed at 52°C. We observed a lower T_sat_ for WT RGG here compared to Fig. 1D due to the need for reduced protein concentration in the case of the shuffled sequences. This finding nicely agrees with our computational results, which showed that increasing the charge segregation in combination with preserving the eIF4E-binding motif enhances self-association propensity more than simply increasing charge segregation. Importantly, despite such drastic rearrangement of the protein sequence, both RGG_shuf_ and RGG_shuf-pres_ formed spherical liquid droplets of normal morphology, as imaged by brightfield microscopy at room temperature (Fig. 2D insets).

To determine whether altering the charge patterning of the RGG sequence has any unexpected consequences in vivo, we then tested RGG_shuf_ in the context of full-length LAF-1 in live yeast cells. LAF-1 in which the RGG domain was replaced with RGG_shuf_ (RGGshuf-LAF-1) appeared to localize to the nucleus, with a single fluorescent punctum per cell (Fig. 2E). This is perhaps unsurprising, as nuclear localization signals characteristically contain stretches of basic amino acids^49, 50^. We, therefore, tagged RGGshuf-LAF1 with a nuclear export signal (NES), which upon expression, generated cytoplasmic puncta, thus demonstrating that RGG_shuf_ is capable of self-assembling in living cells. Together, these experimental results support the computational predictions that charge patterning is a critical determinant of LAF-1 RGG phase separation and that this effect can be supplemented by the incorporation of small patches of hydrophobic amino acids. We were unable to conduct the same experiments on RGG_shuf-pres_ due to its poor expression in yeast cells.

### Arginine and tyrosine are important determinants for LLPS of LAF-1 RGG

Interactions of tyrosine and arginine can be critically important to protein LLPS^30, 32, 51^. The LAF-1 RGG domain contains 24 arginine, 11 tyrosine, and 1 phenylalanine and no lysine residues, which are relatively evenly distributed across the 168 residue-long domain (Fig. 3A). To test the role of these residues in RGG phase separation, in one construct, we mutated all tyrosines to phenylalanine (Y→F), except for a single tyrosine that was mutated to tryptophan to facilitate spectrophotometric detection. In a second construct, we mutated all arginines to lysines (R→K). We then conducted turbidity assays (at 1 mg/mL protein concentration, since the mutations were likely to reduce LLPS propensity) on both constructs.

**Figure 3:**
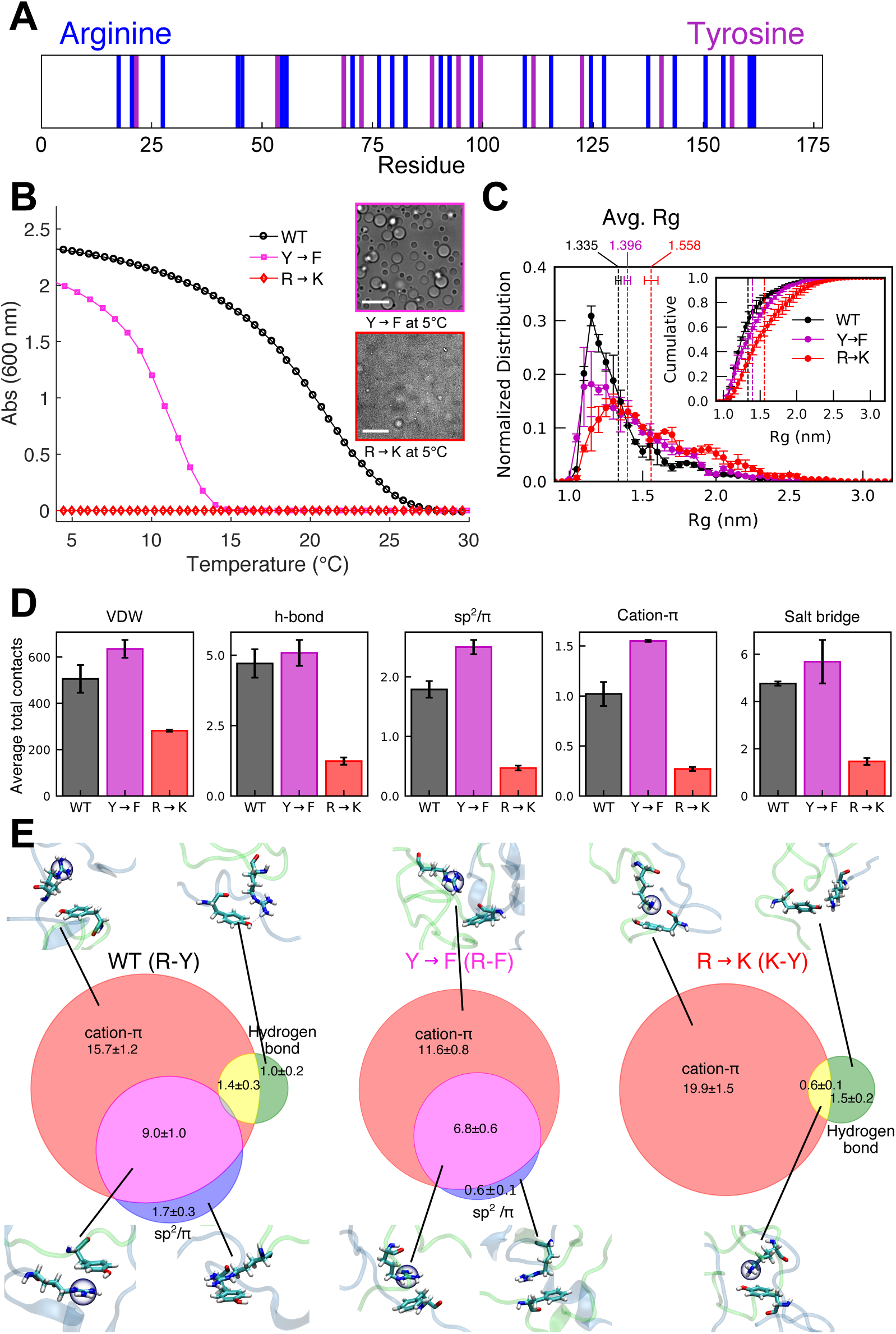
Contribution of arginine and tyrosine residues to LLPS: A) Arrangement of arginine and tyrosine residues along the RGG sequence. Residues are reasonably well-mixed with the exception that the N-terminal end is relatively void of the two amino acids. B) Turbidity measurements show the temperature-dependent phase behavior of WT RGG vs. Y→F or R→K variants. Data shown are representative of three independent turbidity experiments for each protein (Fig. S2). For turbidity assays, protein concentrations were 1 mg/mL (approximately 60 μM) in 150 mM NaCl buffer, pH 7.5. The Y→F variant assembled into spherical liquid droplets (inset micrograph) at 5 °C. The R→K variant did not phase separate in the turbidity assay, nor were micrometer-scale protein liquid droplets visible by optical microscopy (bottom inset), even under conditions favorable for phase separation (6.6 mg/mL protein, 50 mM NaCl, pH 7.5, 5 °C). Scale bars: 10 μm. C) Normalized distribution of radius of gyration (Rg) of RGG**106-149** fragments from single-chain simulations for WT, Y→F, and R→K variants. Inset shows cumulative histogram of Rg. D) Average number of intermolecular contacts observed between two chains of RGG106-149 in two-chain simulations (see Methods), where the average is over the simulated ensemble. Backbone and sidechain heavy atoms are included in these calculations. E) Venn diagrams summarizing the interaction types driving the association of R/K and Y/F residues averaged over all instances of intermolecular VDW contact between any pair of these residues. The numbers represent the percentage and only sidechain heavy atoms are included in these calculations. The overlap between different interaction types shows that they may work cooperatively. WT has all three types of interaction, while R→K loses sp^2^/π interactions, and Y→F loses hydrogen bonding. Snapshots show an instance of indicated contact type(s) from a two-chain simulation. For simulation data, error bars and uncertainty values are SEM with n = 2.

In contrast to WT RGG, which demixed at approximately 26 °C, mutating the tyrosines to phenylalanines lowered transition temperature to approximately 14 °C (Fig. 3B, S2C). To confirm that the Y→F mutant still forms normal protein droplets, we imaged it with brightfield microscopy at 5 °C. We observed that the condensates appeared morphologically identical to WT RGG, with many micrometer-scale protein droplets (Fig. 3B, insets). Even more dramatically, upon mutating all arginines to lysines, we observed no phase separation, even below 5 °C (Fig. 3B, S2C). The R→K mutant was soluble and did not assemble into protein droplets even under experimental conditions that promote RGG phase separation, including high protein concentration and low salt concentration at low temperature. Thus, the presence of tyrosine and arginine plays a key role in phase separation of the LAF-1 RGG domain, in agreement with studies on FUS ^30^.

These experimental results suggest that the Y→F and R→K mutations have a significant impact on the overall interactions occurring between LAF-1 RGG molecules. To gain mechanistic insight into these changes, we turned to all-atom simulations with explicit solvent, which can provide highly detailed information on the different types of interactions in which each amino acid may participate^35^. Since it is currently impractical to faithfully sample the configurational ensemble of a long IDP like LAF-1 using such high-resolution models, we conduct simulations on a 44-residue region of the LAF-1 RGG domain spanning residues 106-149 (RGG106-149). This particular contiguous region was selected to provide the highest compositional similarity with the full RGG domain so that the information obtained is most consistent with the expectations for the full-length sequence (Fig. S5). We also simulate two variants in which either all the tyrosine residues are mutated to phenylalanine (Y→F RGG106-149) or all the arginine residues are mutated to lysine (R→K RGG106-149). From single-chain simulations, we find we find that Rg increases in the following order: WT < Y→F < R→K (Fig. 3C). Previous studies provide compelling evidence that chain dimensions or solvent quality can faithfully provide knowledge on protein LLPS^3, 40, 44, 52^ – more collapsed chains are expected to be more prone to phase separation. Therefore, the trend in Rg from all-atom simulations is consistent with the experimental LLPS behavior that we observe for these mutants (Fig. 3B), which provides further confidence in utilizing these simulations to understand the molecular interactions responsible for the experimental results.

To observe intermolecular interactions and self-association, we conducted simulations of two RGG106-149 chains. Consistent with our recent work on the FUS LC domain^35^, we use well-tempered metadynamics with the number of intermolecular Van der Waals (VDW) contacts as a pertinent collective variable to enhance sampling of intermolecular contacts between the two peptides. The resulting free energy surfaces as a function of the number of intermolecular VDW contacts are shown in Fig. S5A. Both WT and Y→F peptides show free energy minima at a finite number of VDW contacts. Interestingly, the R→K variant has a global minimum at zero contacts, suggesting the two chains do not interact as is consistent with the lack of phase separation in the experiments.

Previous work has suggested the importance of cation-π interactions^30, 53^, particularly between arginine and tyrosine^30^; planar interactions between sp^2^ hybridized groups (referred to here as sp^2^/π interactions)^54^; electrostatic interactions^16, 55^; and hydrophobic and VDW interactions^35^ to LLPS. We calculate the average number of intermolecular contacts between the two chains of the different RGG106-149 variants (Fig. 3D). In general, WT and Y→F have a much higher number of contacts than R→K, consistent with the free energy profiles, showing that R→K most favors unbound configurations. We also normalize the average number of intermolecular contacts of each type by the average number of intermolecular VDW contacts (Fig. S6B) to understand the role of various interaction modes independent of the global contact propensity, which is different between these three variants. Additionally, we provide the unnormalized average number of various contacts formed by each residue (Fig. S7). The number of sp2/pi and cation-pi interactions is particularly decreased in R→K, while there is no significant difference between WT and Y→F average contacts. The overall number of contacts, however, may not consider the interaction strengths and thus would not perfectly describe the difference between WT and Y→F.

To further elucidate the differences between the mutants, we considered the effect of the interactions between cationic and aromatic sidechains which were the original target of these designed mutations. By analysis of all simulation snapshots in which arginine or lysine and tyrosine or phenylalanine residues from different chains are in contact (having at least one VDW contact between them), we calculated the probability of occurrence of different interaction types. Three different interaction modes are observed for arginine-tyrosine contacts, while only two are observed for arginine-phenylalanine and lysine-tyrosine contacts (Fig. 3E). Importantly, interactions between arginine-tyrosine sidechains promoting LLPS could be due to multiple modes of interactions with significant contributions from cation-π, hydrogen bonding, and sp^2^/π interactions. The Y→F mutations reduce the extent of these interactions, likely due to the loss of hydrogen bonding interactions, as phenylalanine sidechain lacks a hydroxyl group, unlike tyrosine. On the other hand, R→K mutations remove sp^2^/π interactions due to the removal of the guanidine group present in arginine. These results provide a much-needed mechanistic understanding of the importance of arginine and tyrosine residues to protein LLPS.

### Sequence perturbations result in shifts to phase diagram

To more completely map the experimental phase behavior of variants of the LAF-1 RGG domain, we performed temperature-dependent turbidimetry at varying protein concentrations and calculated T_sat_ for each to obtain the low-concentration arm of their phase diagrams. We find that all variants for which we were able to acquire multiple T_sat_ values display a UCST phase diagram, having a region of miscibility at high temperatures and phase separation at low temperatures. By imposing different perturbations to the RGG sequence, we were able to shift the phase diagram upward (Fig. 4A) or downward (Fig. 4B). A significant increase of LLPS propensity occurs when modifying the sequence such that most cations are localized to one side and anions on the other side, even when the sticky hydrophobic region we identified is lost in the shuffling. We find that designing a shuffled sequence that conserves this region (such conservation has occurred across different organisms) results in the greatest upward shift of the phase diagram (Fig. 4A), indicating that both of these types of molecular interactions control phase separation of RGG.

**Figure 4:**
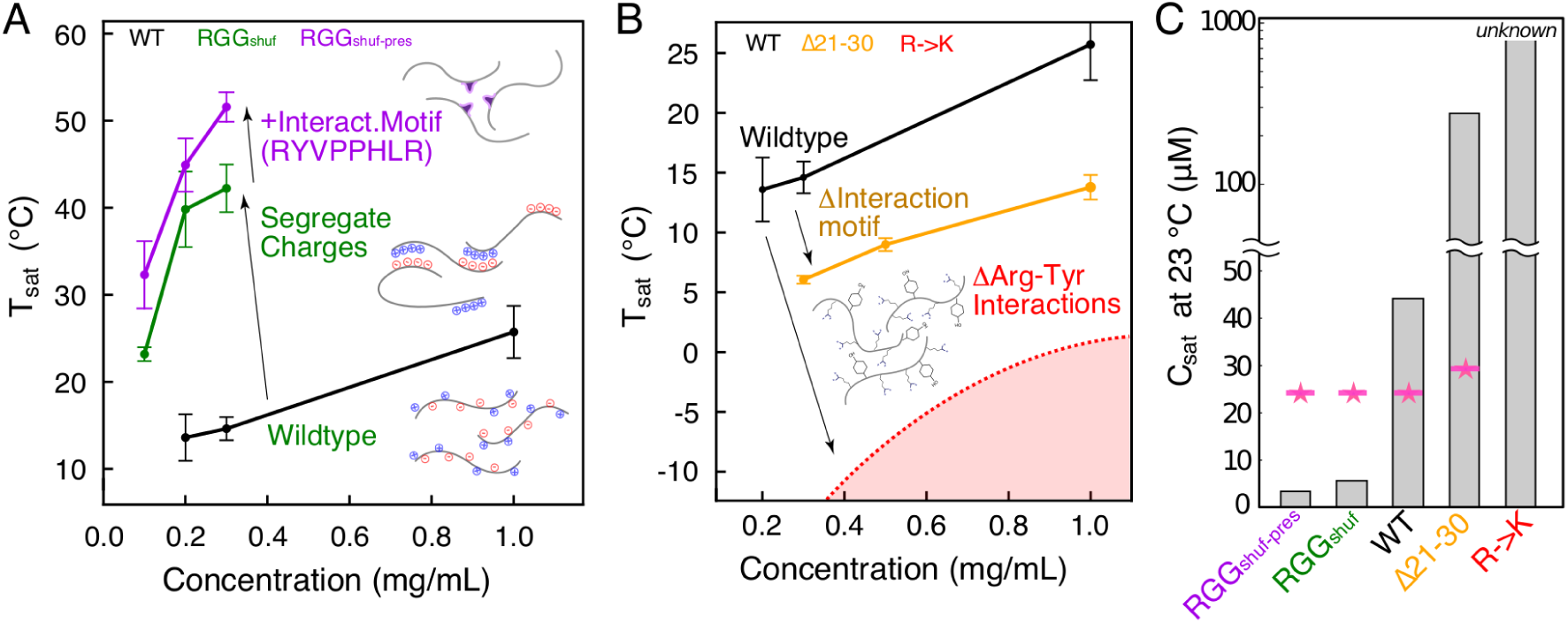
Phase diagrams illustrate molecular interactions that underlie RGG LLPS: Phase diagrams for different LAF-1 variants. T_sat_ values and associated error bars were calculated from triplicates of the turbidity assays at each concentration. A) Shuffled sequences with a high degree of charge patterning shift phase diagram upward, making phase separation occur at lower concentrations more easily. RGG_shuf-pres_ features both charge segregation and the self-interaction motif at residues 21-28, allowing for even greater LLPS propensity. B) Deletion of the interaction motif, or mutation of arginine residues to lysine, both result in a drastic decrease of LLPS propensity and downward shift of the phase diagram. Phase diagram for R→K is theoretical and is meant strictly as a visual guide to show that this mutation has a stronger effect on LLPS than the deletion of the interaction motif. T_sat_ of WT, Δ21-30, RGG_shuf_, and RGG_shuf-pres_ are all significantly different than one another (p < 0.005), based on one-way ANOVA followed by Tukey’s post-hoc test at 0.3 mg/mL. C) Saturation concentrations from turbidity experiments compared with predictions presented in ref ^30^.

We are also able to shift the phase diagram downward, thus making LLPS less favorable. When deleting residues 21-30, encompassing the eIF4E-binding motif, we find that the phase diagram shifts downward significantly (Fig. 4B), much more so than when deleting other regions of 10 residues (Fig. S8). This also validates the predictions of the computational model, which identified the enhanced interactions within that region. Mutations of all arginine to lysine result in total loss of LLPS behavior at tested conditions. We suggest the phase diagram has been shifted downward enough that the temperatures or concentrations required to observe LLPS are not practically achievable in vitro (Fig. 4B).

In previous work, Wang et al. suggest that the saturation concentration (c_sat_) of a protein may be predicted by counting the number of tyrosine and arginine residues within the sequence as *c*_*sat*_ = *k*(*n*_*Tyr*_*n*_*Arg*_)^−1^. where k is a fitting parameter and is equal to 6.5 mM^30^. For the WT RGG sequence, this predicts a saturation concentration of 24.6 μM or 0.439 mg/mL, which also applies to RGG_shuf_ and RGG_shuf-pres_, as they have an identical composition (Fig. 4C). For the R→K and Y→F variants, the denominator becomes zero, so the predicted value is undefined, with the suggestion that c_sat_ is very high. Deletion of residues 21-30 removes 2 arginine, and 1 tyrosine residue, resulting in a small predicted increase of c_sat_ to29.6 μM or 0.493 mg/mL. To directly compare with results from this prediction, we calculated saturation concentration at 23°C using a logarithmic fit to turbidimetry data (Fig. S9A,B). Linear fits of the data yield similar c_sat_ values (Fig. S9C,D). We find that the equation *c*_*sat*_ = *k*(*n*_*Tyr*_*n*_*Arg*_)^−1^ poorly predicts the c_sat_ for RGG_shuf_ or RGG_shuf-pres_ (Fig 4C). Further, the prediction underestimates the effect of deletion of residues 21-30 from RGG. These results suggest that while the number of arginine and tyrosine residues can sometimes provide a reasonable estimate of c_sat_, this parameter alone is not predictive, and many other factors, such as charge patterning and hydrophobic interactions, determine LLPS.

### Protein condensates formed from RGG variants retain liquid-like properties

Thus far, we have demonstrated perturbations to the LAF-1 RGG sequence that alter its phase behavior, using molecular simulations to guide experiments and provide a mechanistic understanding of the driving forces of phase separation. We next wondered whether these sequence perturbations would alter the liquid properties of RGG protein condensates. This is important to understand because the material properties of biomolecular condensates are intertwined with their biological function^56^. The spherical morphologies of WT RGG and its sequence variants are characteristic of viscous liquids. For all variants, droplets could be seen contacting, fusing, and then rounding into larger spheres (Fig. 5A). To determine the liquidity of these droplets, we quantified fusion events, calculating the time τ for the two coalescing droplets to relax to a sphere (Fig. S10A). WT RGG and all the variants examined (RGGΔ21-30, both shuffled versions, and Y→F) exhibited rapid fusion, with droplets of lengthscale ℓ = 2 ± 0.25 μm fusing with τ < 100 ms (Fig. 5B). Droplet fusion is driven by surface tension γ and slowed by viscosity η, and the timescale of fusion is also proportional to droplet size ℓ, so 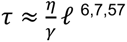. By plotting τ against ℓ for tens of droplet fusion events (Fig. 5C), we estimate the ratio η/γ, known as the inverse capillary velocity (Fig. S10B). All the variants tested had η/γ within 3-fold that of WT RGG, and in all cases η/γ < 0.05 s/μm, indicating faster fusion compared to LAF-1 (η/γ = 0.12 s/μm)^6^.

**Figure 5:**
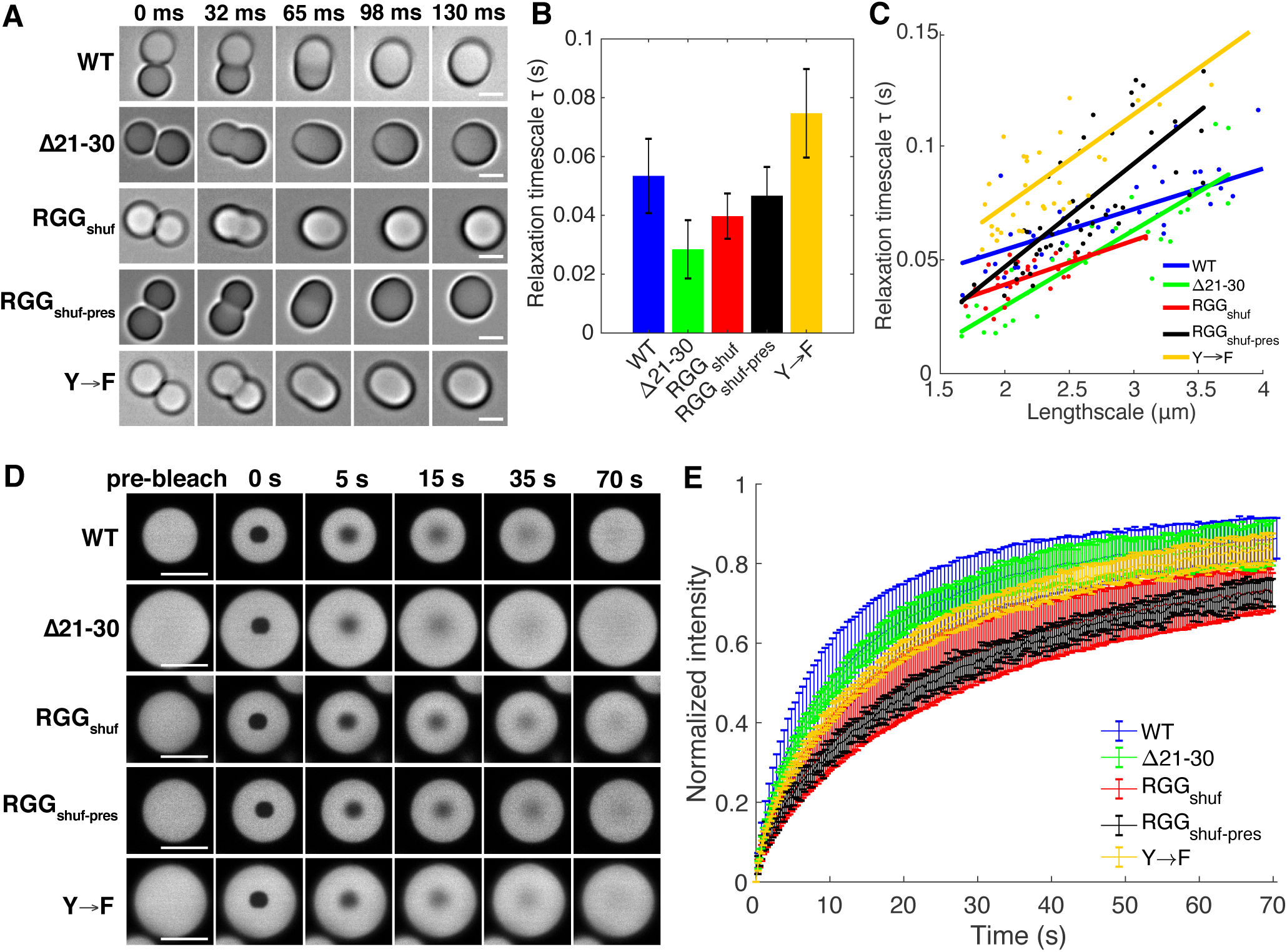
RGG variants exhibit liquid-like material properties. (A) Droplets fuse rapidly to form a single larger sphere. Scale bars: 2 μm. (B) The timescale of droplet fusion for droplets of lengthscale range 1.75-2.25 μm. Error bars represent STD (n ≥ 9). (C) Plots of relaxation timescale against length scale, which allows determination of the inverse capillary velocity. (D) Representative images from FRAP experiments. Scale bars: 5 μm. (E) Normalized FRAP recovery curves show > 50% recovery within 30 s for all variants. Error bars represent STD (n > 15).

In a complementary approach, we examined dynamics within the droplets through fluorescence recovery after photobleaching (FRAP). For all variants tested, 50% fluorescence recovery was achieved within 30 s of photobleaching a small circular region within a larger droplet (Fig. 5D,E). By fitting the FRAP recovery curves to a 3D infinite model, we find diffusion coefficients ranging from D = 0.01 μm^2^/s to 0.025 μm^2^/s, approximately one order of magnitude faster than that for full-length LAF-1^58^. There are modest differences, notably that the construct with deletion of residues 21-30 (ie lower T_sat_ that WT) exhibited faster fusion and FRAP recovery compared to that of RGGshuff-pres (ie highest T_sat_ of all constructs we tested). Together, the FRAP and fusion experiments demonstrate that these variants maintain dynamic, liquid-like condensates, despite the changes to sequence and phase behavior. Thus, phase behavior – critical concentration and transition temperature – can be modulated mostly independently from the alteration of droplet liquidity.

## Discussion

In this work, we elucidate sequence determinants of IDP phase separation, and in so doing, we advance a computationally guided approach for rational engineering of protein LLPS. We focus on the RGG domain from LAF-1, a prototypical phase-separating protein of great interest to the LLPS field whose sequence-to-phase behavior relationship has not been mapped in detail previously. By combining predictive simulations and experiments, we identified three important features that govern the propensity of this protein to phase separate: a short conserved domain, charge patterning, and arginine-tyrosine interactions.

We first demonstrate that a small conserved domain plays an unexpectedly large role in LAF-1 phase separation, such that the deletion of 10 residues encompassing the identified region decreases the protein’s phase separation propensity significantly. Our computational data and in vitro experiments show that this region has an intrinsic affinity for itself. This contact-prone region coincides with the previously identified eIF4E-binding motif, although the contribution of this motif to LLPS is likely orthogonal to its specific binding function. Hypothetically, LLPS of LAF-1 might be particularly sensitive to stimuli that may target this region, such as phosphorylation-induced folding that may hide the motif and block its accessibility for self-association^59^. More generally, these results suggest that the presence within proteins of functional motifs, such as specific binding motifs^38^, may have a non-negligible effect on LLPS of the full sequence – even if the functional motif is only a small region in a much larger protein.

Second, our results support a revised view of the role of electrostatic interactions in LAF-1 RGG phase separation. Previous views pointed to electrostatic interactions and charge patterning as the driving force for LAF-1 phase separation^6, 13^. On the contrary, we found that WT LAF-1 RGG has a well-mixed charge distribution. We, therefore, asked whether introducing charge patterning could enhance LAF-1 phase separation. We used the SCD metric to identify shuffled versions of LAF-1 RGG having a high degree of charge segregation, and our CG simulations and experiments both show that such charge patterning results in significantly enhanced propensity to phase separate. Our results extend previous work on this topic^44, 45, 60^. Ddx4 features blocks of alternating net charge, and scrambling the blocks to remove charge patterning abolishes phase separation^47^. Relatedly, complex coacervation of the negatively charged Nephrin intracellular domain (NICD) with positively charged partners is promoted in part by blocks of high charge density in NICD^55^. Theoretical work shows as well that block polyampholytes exhibit stronger interactions compared to charge-scattered polyampholytes, as the latter experience repulsion from nearby like charges^44^. Thus, it appears that WT RGG may be under negative selection to moderate this mode of blocky electrostatic interaction and maintain a well-mixed charge distribution.

Third, we find that distributed tyrosine and arginine residues are also important to the ability of LAF-1 RGG to phase separate, and we gain valuable mechanistic insight into this result from all-atom simulations. The importance of these particular residues was attributed in previous work to their propensity to form cation-π interactions^30, 32, 61^. Our all-atom simulations confirm the presence of cation-π interactions and, importantly, highlight other important interaction modes as well that change when mutating arginine to lysine or tyrosine to phenylalanine. Our simulations suggest that the loss of planar sp^2^/π interaction^54^ is likely responsible for reduced LLPS when mutating arginine to lysine. We note that arginine may be particularly prone to promoting LLPS with aromatic-rich sequences due to cooperative cation-π and sp^2^/π interactions that co-occur. Another important interaction mode is hydrogen bonding, which has also recently been demonstrated to be important to LLPS^35, 37^ and is present in interactions between cationic residues and tyrosine. Our simulations suggest that the reduced LLPS propensity when mutating tyrosine to phenylalanine can be explained by the loss of sidechain hydrogen bonding, as phenylalanine lacks the hydroxyl group. Therefore, we suggest that while the selected mutations likely weaken cation-π interactions^30, 32^, one must also consider the loss of several other types of interactions that are responsible for stabilizing the condensed liquid phase^35^.

The sequence perturbations investigated here significantly altered c_sat_ – for instance, approximately one order of magnitude decrease in c_sat_ for RGGshuff-pres compared to WT RGG, and approximately 5-fold increase for Δ21-30. Remarkably, we observed that the RGG variants retained their dynamic liquid material properties, even for a perturbation as drastic as shuffling the sequence. The significant changes in phase behavior would likely have important biological consequences, whereas the modest differences in droplet fluidity are likely of smaller functional significance. Thus, our experiments suggest that in a predictive manner, we can design mutations to an IDP to alter its phase behavior while retaining liquid-like condensate dynamics. Future work will continue to explore sequence-to-rheology relationships^30^.

Overall, our combined results elucidate the driving forces of LLPS and highlight how the sequence perturbations affect LLPS, promising a framework toward the rational design of LLPS-enabled IDPs. This work will inform studies into the biology of membraneless organelles, aberrant phase transitions in disease, and design of biomaterials and synthetic organelles.

## Methods

### Cloning

The WT, full-length LAF-1 gene was a gift of Shana Elbaum-Garfinkle and Clifford Brangwynne. WT RGG was amplified by PCR from LAF-1. All modified versions of the RGG domain were ordered as synthetic double-stranded DNA fragments (gBlocks; IDT). Plasmids were constructed using either In-Fusion cloning (Takara Bio) or NEBuilder HiFi DNA Assembly (New England BioLabs). For bacterial expression, genes were cloned into a pET vector in-frame with a C-terminal 6xHis-tag. For yeast expression, genes were cloned into the YIplac211 vector in frame with a C-terminal mEGFP (monomeric enhanced GFP) tag. YIplac211 is a yeast integrating plasmid with a URA3 marker^62^. Gene sequences were verified by Sanger sequencing (GENEWIZ).

### Protein expression and purification

For bacterial expression, plasmids were transformed into BL21(DE3) competent *E. coli* (New England BioLabs). Colonies picked from fresh plates were grown for 8 h at 37 °C in 1 mL LB + 1% glucose while shaking at 250 rpm. This starter culture (0.5 mL) was then used to inoculate 0.5 L cultures. Cultures were grown overnight in 2L baffled flasks in Terrific Broth auto-induction medium (Formedium; supplemented with 4 g/L glycerol) at 37 °C while shaking at 250 rpm. The pET vectors used contained a kanamycin resistance gene; kanamycin was used at concentrations of 50 μg/mL in starter cultures and 100 μg/mL in the auto-induction medium^63^. After overnight expression, bacterial cells were pelleted by centrifugation. Pellets were resuspended in lysis buffer (1 M NaCl, 20 mM Tris, 20 mM imidazole, Roche EDTA-free protease inhibitor, pH 7.5) and lysed by sonication. Lysate was clarified by centrifugation at 15,000 g for 30-60 minutes. Lysis was conducted on ice, but other steps were conducted at room temperature to prevent phase separation. Proteins were purified using an AKTA FPLC with 1 mL nickel-charged HisTrap columns (GE Healthcare Life Sciences) for affinity chromatography of the His-tagged proteins. The column was washed with 500 mM NaCl, 20 mM Tris, 20 mM imidazole, pH 7.5. Proteins were eluted with a linear gradient up to 500 mM NaCl, 20 mM Tris, 500 mM imidazole, pH 7.5. Proteins were dialyzed overnight using 7 kDa MWCO membranes (Slide-A-Lyzer G2, Thermo Fisher) into 500 mM NaCl, 20 mM Tris, pH 7.5 or 150 mM NaCl, 20 mM Tris, pH 7.5. Proteins were dialyzed at temperatures (25 °C -42 °C) high enough to inhibit phase separation because phase-separated protein bound irreversibly to the dialysis membrane. Proteins were snap frozen in liquid N2 in single-use aliquots and stored at -80 °C. For turbidity and microscopy experiments, protein samples were prepared as follows: Protein aliquots were thawed above the phase transition temperature. Proteins were then mixed with buffer (20 mM Tris, pH 7.5, 0 – 150 mM NaCl) to obtain solutions containing the desired protein and NaCl concentrations. Protein concentrations were measured based on their absorbance at 280 nm using a Nanodrop spectrophotometer (ThermoFisher). Proteins were mixed in a 1:1 ratio with 8 M urea to prevent phase separation during concentration measurements.

### MALDI-TOF mass spectrometry

Molecular weights of purified proteins were measured by matrix-assisted laser desorption/ionization time-of-flight (MALDI-TOF) mass spectrometry on an Ultraflextreme mass spectrometer (Bruker). Protein samples were applied as spots to an MPT 384 polished steel target plate. Spots consisted of 1 μL protein solution (approximately 10 μM protein in 50 mM NaCl) plus 1 μL matrix solution (10 mg/mL sinapinic acid dissolved in a 50:50 acetonitrile:water mixture with 0.1% trifluoroacetic acid added).

### Turbidity assays

Temperature-dependent turbidity assays were conducted using a UV-Vis spectrophotometer (Cary 100 Bio; Agilent) equipped with a multicell Peltier temperature controller. Protein samples were assayed in quartz cuvettes with 1 cm path length (Thorlabs). Samples were first equilibrated above the phase transition temperature (25-60 °C depending on the sample) and blanked. Then, the samples were cooled at a rate of 1 °C per minute until reaching 2 °C. Absorbance was measured at λ = 600 nm every 0.5 °C throughout the temperature ramp. Upon cooling below the phase transition temperature, the samples changed from clear to turbid.

### SDS-PAGE and western blot

For chromatographically purified proteins, SDS-PAGE was run using NuPAGE 4-12% Bis-Tris gels (Invitrogen) and stained using a Coomassie stain (SimplyBlue SafeStain; Invitrogen). For western blotting, yeast cells were lysed as follows^64^: Cell cultures were pretreated with 2 M lithium acetate for 5 minutes on ice, then with 0.4 M NaOH for 5 minutes on ice. The cell cultures were then resuspended in SDS sample buffer, heated at 95 °C for 5 minutes, and centrifuged to remove cell debris. The supernatant was stored at -80 °C until use. The supernatant was run on a Novex 10% Tris-Glycine gel, WedgeWell format (Invitrogen), then transferred to a nitrocellulose membrane (0.2 μm pore size). The membrane was then incubated with two primary antibodies: rabbit polyclonal antibody to GFP (Invitrogen, catalog #A11122) for detection of the GFP-tagged LAF-1 constructs, and mouse monoclonal antibody to PGK1 (Invitrogen, catalog #459250) as a loading control. Secondary antibodies used for detection were IRDye 680RD goat anti-rabbit IgG (LI-COR, catalog #926-68071) and IRDye 800CW goat anti-mouse IgG (LI-COR, catalog #926-32210). Blots were visualized on a LI-COR Odyssey CLx infrared imaging system.

### Yeast transformation and yeast cultures

YIplac211 plasmids were prepared for yeast chromosomal integration by restriction digest with EcoRV, which cuts in the URA3 marker. Linearized plasmids were transformed into *S. cerevisiae* YEF473A strain^65^ using the Frozen-EZ Yeast Transformation II Kit (Zymo Research). Transformed yeast cells were cultured at 30°C in uracil dropout synthetic defined medium (-Ura dropout supplement was purchased from Takara Bio). To induce expression of genes under the control of the GAL1 promoter, yeast cultures were first grown overnight in dropout medium + 2% glucose, then grown for 8-10 hours in dropout medium + 2% raffinose, and finally grown overnight in dropout medium + 2% galactose with a target OD600 = 0.3 – 0.5 for imaging.

### Microscopy: phase behavior, FRAP, and fusion

Imaging of temperature-dependent phase behavior in vitro and in yeast was performed on an Olympus IX81 inverted microscope equipped with a Yokogawa CSU-X1 spinning disk confocal unit and an iXon3 EMCCD camera (Andor). The microscope stage was outfitted with a Cherry Temp microfluidic temperature controller (Cherry Biotech), which enabled imaging samples over the temperature range 5 to 42 °C, with rapid switching (approximately 10 s) between temperature extremes. Imaging was conducted with a 100x/1.4 NA plan-apochromatic oil-immersion objective.

FRAP experiments were performed on a Zeiss Axio Observer 7 inverted microscope equipped with an LSM900 laser scanning confocal module and a 63x/1.4 NA plan-apochromatic oil-immersion objective. LAF-1 RGG and its variants were mixed with 5% of RGG-GFP-RGG, which partitions into the RGG droplets and serves as a FRAP probe^21^. GFP was imaged with a 488 nm laser and bleached with a 405 nm laser. Circular bleach regions of approximate radius R = 1.5 μm were drawn in the center of protein droplets whose radii were at least 2.5R. Recovery curves were fit to an infinite boundary model in three dimensions to calculate the recovery timescale τ^58^. The diffusion coefficient was calculated as D = R^2^/τ. The same Zeiss microscope was used for droplet fusion experiments, but using brightfield transillumination and imaging onto an Axiocam 702 sCMOS camera at a frame rate of approximately 62 Hz. Droplet fusion was analyzed by first fitting the image of the fusing droplets to an ellipse and calculating the aspect ratio of the ellipse. The aspect ratio was then plotted against time and the decreasing portion of the curve was fit to an exponential decay to calculate the relaxation time^7, 27, 57^. The droplet length scale was defined as the radius of the droplet after completion of fusion, when the merged droplet was circular (aspect ratio 1). FRAP and droplet fusion experiments were conducted at room temperature of 16-18 °C using protein concentrations above c_sat_ at that temperature. Image analysis and data processing were performed in MATLAB.

All other imaging was performed on a Leica DMi8 inverted microscope equipped with a spinning disk confocal unit (Spectral Applied Research) and an sCMOS camera (Orca Flash 4.0; Hamamatsu) using a 63x/1.4 NA or 100x/1.4 NA plan-apochromatic oil-immersion objective.

For imaging purified RGG proteins, the protein samples were placed in chambers on glass coverslips (#1.5 glass thickness) that had been passivated for >1 hr by incubation with 5% Pluronic F127 (for FRAP and droplet fusion experiments) or bovine serum albumin. Coated coverslips were thoroughly rinsed with buffer prior to the addition of RGG protein solutions. For imaging yeast, the glass surface was pretreated by incubation with 0.4 mg/mL concanavalin A (ConA; Sigma) for 5-10 minutes. After removing the ConA solution, yeast was pipetted into the imaging chamber and allowed to settle for several minutes before imaging.

### Coarse-grained simulations

Coarse-grained simulations were conducted using an amino-acid-resolution model with 20 residue types to capture sequence specificity, having interactions based on relative hydropathies of each amino acid. Each system was simulated at a range of temperatures using constant volume and temperature using a Langevin thermostat, following similar protocols to our previous work^26^. Simulations of phase coexistence were conducted using HOOMD-Blue v2.1.5 software package^66^.

### All-atom simulations

Atomic-resolution simulations were conducted for systems containing either one or two copies of a 44-residue fragment of the LAF-1 RGG domain (RGG106-149). Simulations were of 44-residue fragments as we have found this size to be computationally tractable for single- and two-chain simulations in the previous studies^5, 35^. We selected residues 106-149 by calculating the overall sequence composition of all possible 44-residue fragments and comparing them with the total composition of the 168-residue RGG domain (Fig. S5A). The region having the overall composition most similar to that of the full RGG domain was residues 106-149. Notably, this fragment contains 6 arginine and 3 tyrosine residues constituting 13.6% and 6.8% of the 44-residue sequence, comparable to the 14.3% and 6.5% composition in the full RGG (Fig. S5B,C).

Simulations were conducted with either a single RGG106-149 chain solvated in explicit water and ∼100 mM NaCl or two chains at the same conditions. We used a modified version of the state-of-the-art Amber99SBws force field^67^ with improved residue-specific dihedral corrections (unpublished), tip4p/2005 water^68^ and improved salt parameters from Luo and Roux^69^. To efficiently sample the configurational ensemble and contacts between amino acid residues, we employed enhanced sampling using parallel tempering in the well-tempered ensemble (PT-WTE) which couples replica exchange molecular dynamics (REMD)^70^ and well-tempered metadynamics^71^ applied to the total system energy to enhance fluctuations and reduces the number of replicas required for good replica exchanges^72^. For two-chain simulations, we also applied a well-tempered metadynamics bias on the intermolecular VDW contacts between heavy nonpolar atoms (i.e. |q| < 0.25) as we have done previously to improve sampling of binding and unbinding events^35^. Simulations were conducted using GROMACS 2016 software package^73^ with PLUMED 2.4 plugin^74^.

We calculated the free energy surface of the two-chain systems from the metadynamics bias using the built-in function (sum_hills) in PLUMED, and an alternative time-independent method from Tiwary and Parrinello^75^, then subtract the difference between the two results to generate error bars for Fig. S6A. Contact propensities in all-atom two-chain PT-WTE simulations were reweighted based on free energy surface.

VDW contacts were considered as any two heavy atoms being within 6 Å of each other. Hydrogen bonds were considered as a donor atom and an acceptor atom being within 3 Å and the donor-hydrogen-acceptor angle being larger than 120°. Sp^2^/π interactions were calculated as presented by Vernon et al.^54^ and considered as any two sp^2^-hybridized groups having at least two pairs of atoms being within 4.9 Å and the angle between the normal axes of the two sp2-planes being less than 60°. Cation-π interactions were considered as a cationic atom being within 7 Å of the center of an aromatic ring and less than 60° from the normal axis of the π face. Salt bridges are considered as a cationic atom and an anionic atom being within 6 Å of each other.

## Author Contributions

BSS, GLD, MCG, and JM conceived and designed research. BSS, FMK, AKR, CNJ, and AGS conducted experiments. GLD, WST, and RMR performed and analyzed simulations. BSS, DAH, MCG, and JM supervised research. BSS, GLD, and JM wrote manuscript.

## Acknowledgements

We thank Erfei Bi, Kangji Wang, and James Shorter for yeast strains, reagents, and protocols, and Cliff Brangwynne and Shana Elbaum-Garfinkle for the full-length LAF-1 gene. We gratefully acknowledge Andrew Tsourkas for use of the temperature-controlled spectrophotometer, Hui Chen for assistance with western blotting, Ellen Reed for assistance with mass spectrometry, Xinyi Li for assistance with data analysis, and Nick Fawzi for helpful discussions. This work was supported by the U.S. Department of Energy, Office of Science, Basic Energy Sciences awards DE-SC0007063 to D.H. (experiments) and DE-SC0013979 to J.M. (theory and simulation). We gratefully acknowledge the use of the high-performance computing capabilities of the Extreme Science and Engineering Discovery Environment (XSEDE), which is supported by NSF grant TG-MCB-120014, and the National Energy Research Scientific Computing Center, supported by the Office of Science of the U.S. Department of Energy under contract DE-AC02-05CH11231. B.S. received support from an NIH postdoctoral fellowship (F32GM119430). W.S.T. received support from a National Science Foundation Grant (1845734). M.G. acknowledges support from a National Science Foundation Superseed, NIH R01-EB028320, and Burroughs Wellcome Fund.

## 1. Supporting Text

### 1.1 **Sequences used in in vitro work** (including His tag and XhoI restriction site)

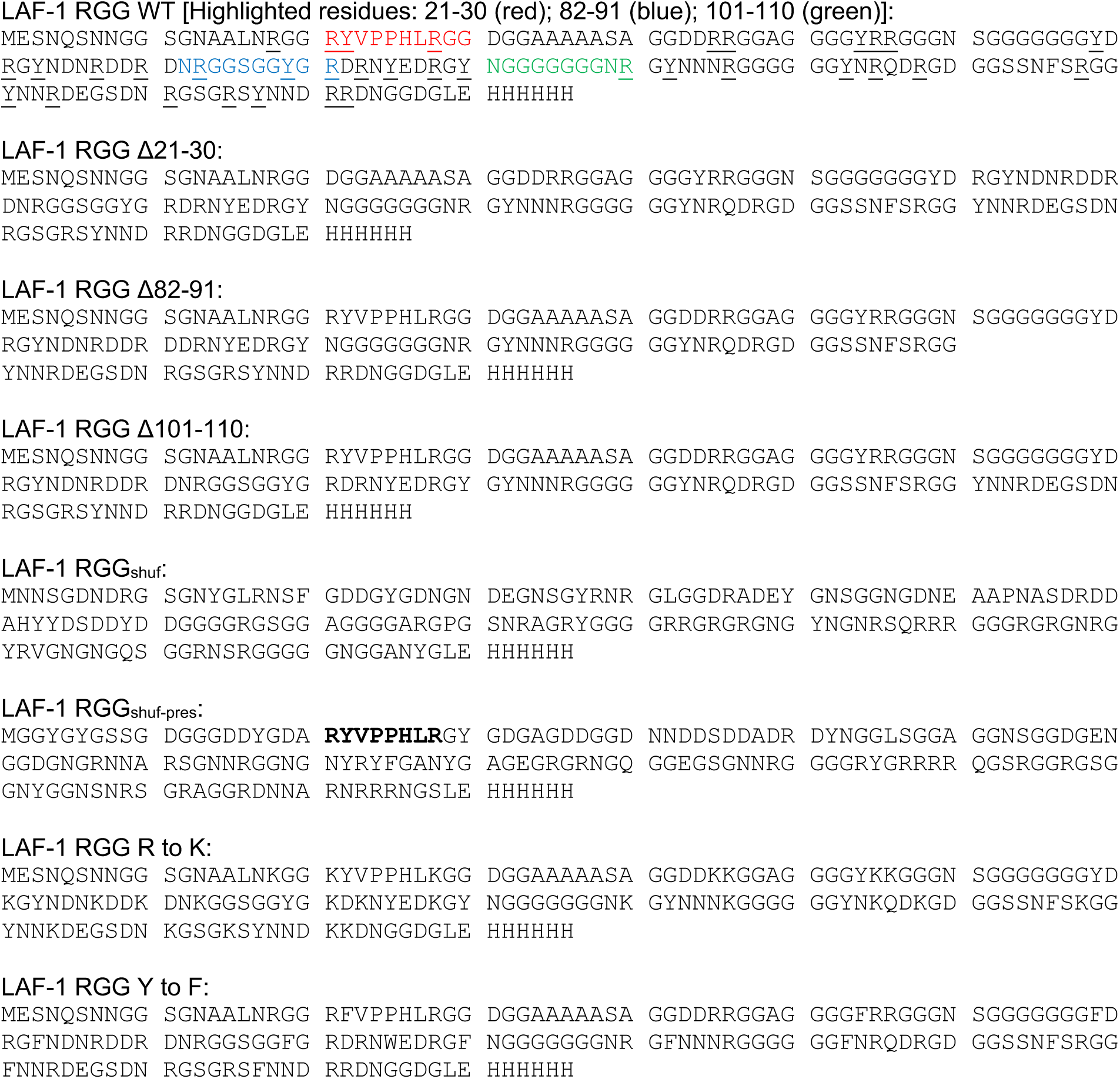

### 1.2 LAF-1 homologs used in sequence alignment (accession numbers)

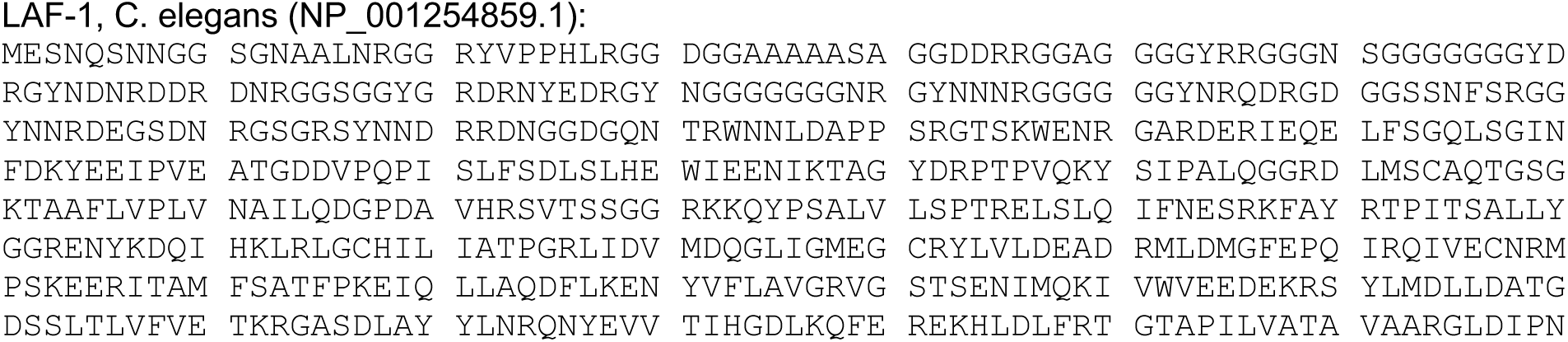
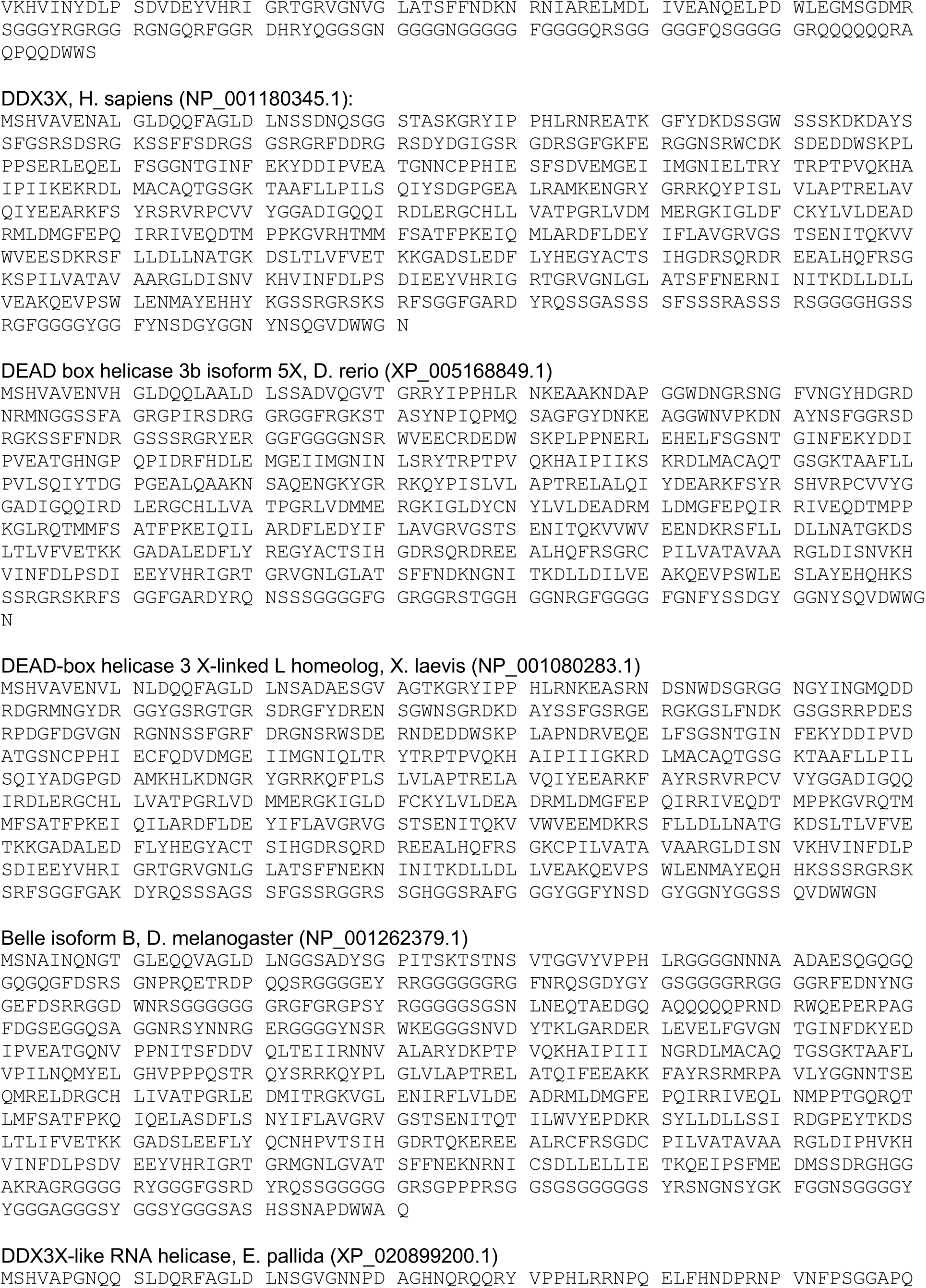
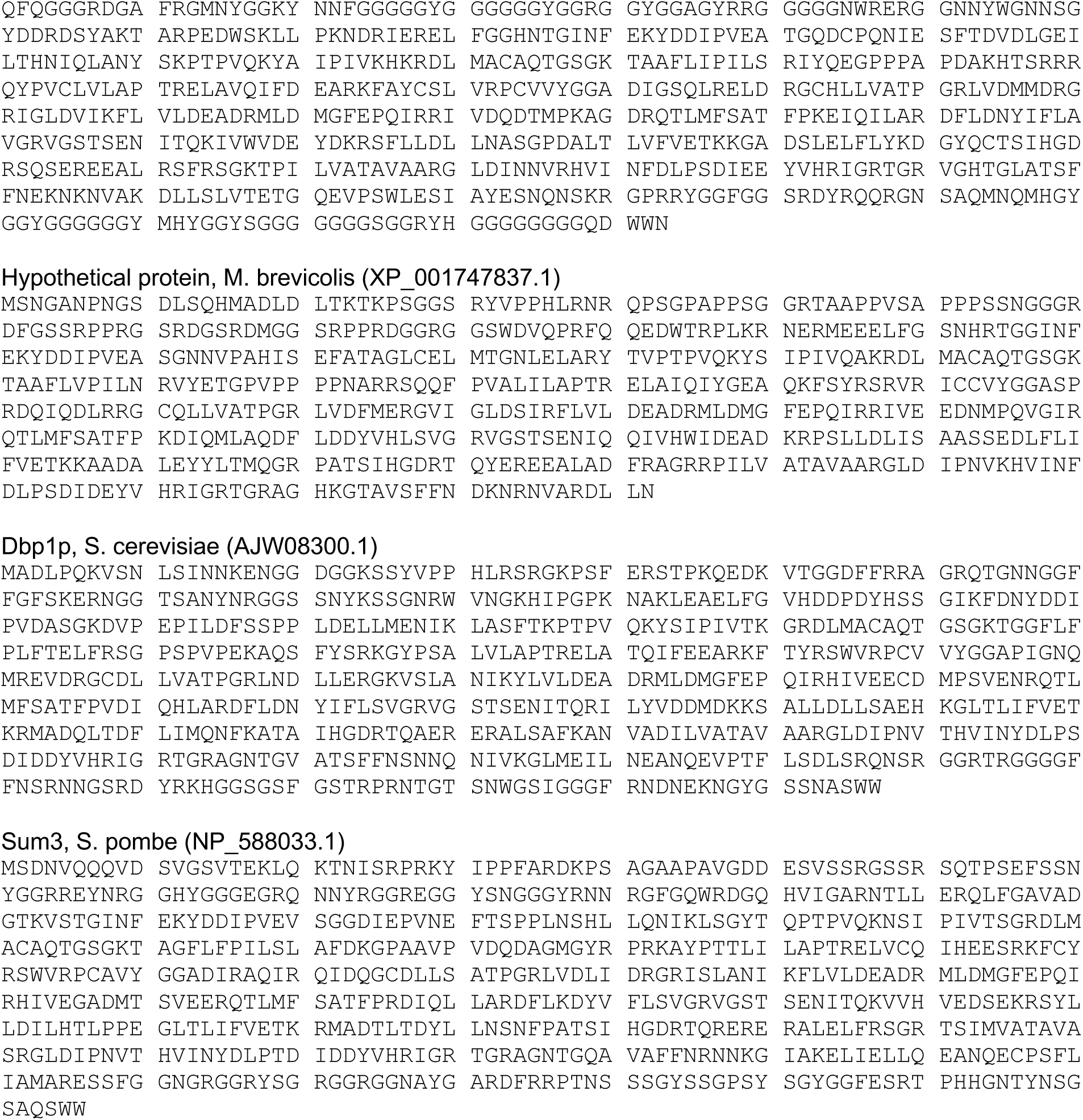

### 1.3 Homolog sequence alignment

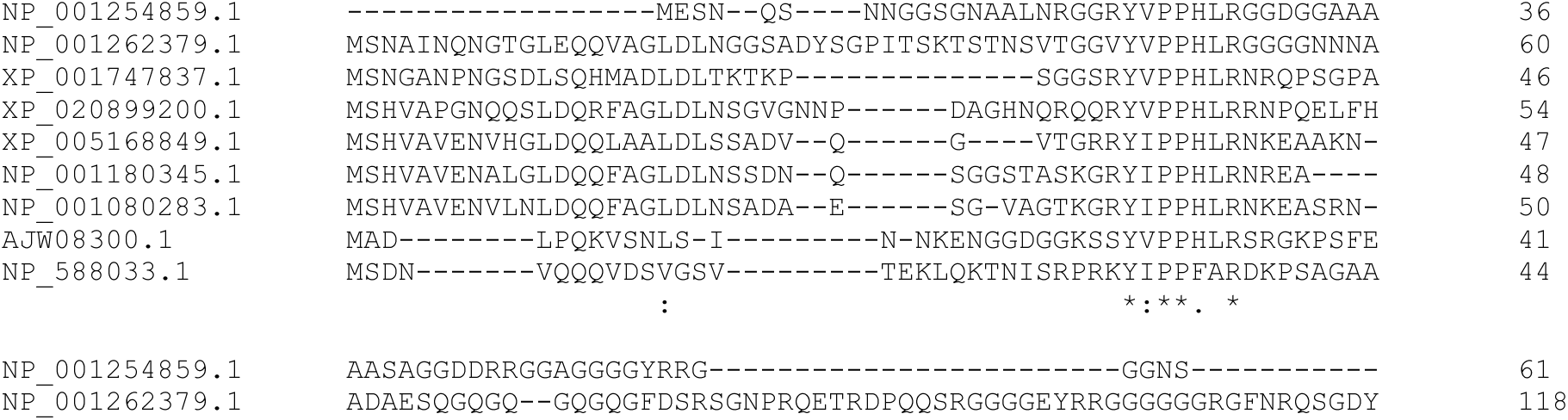
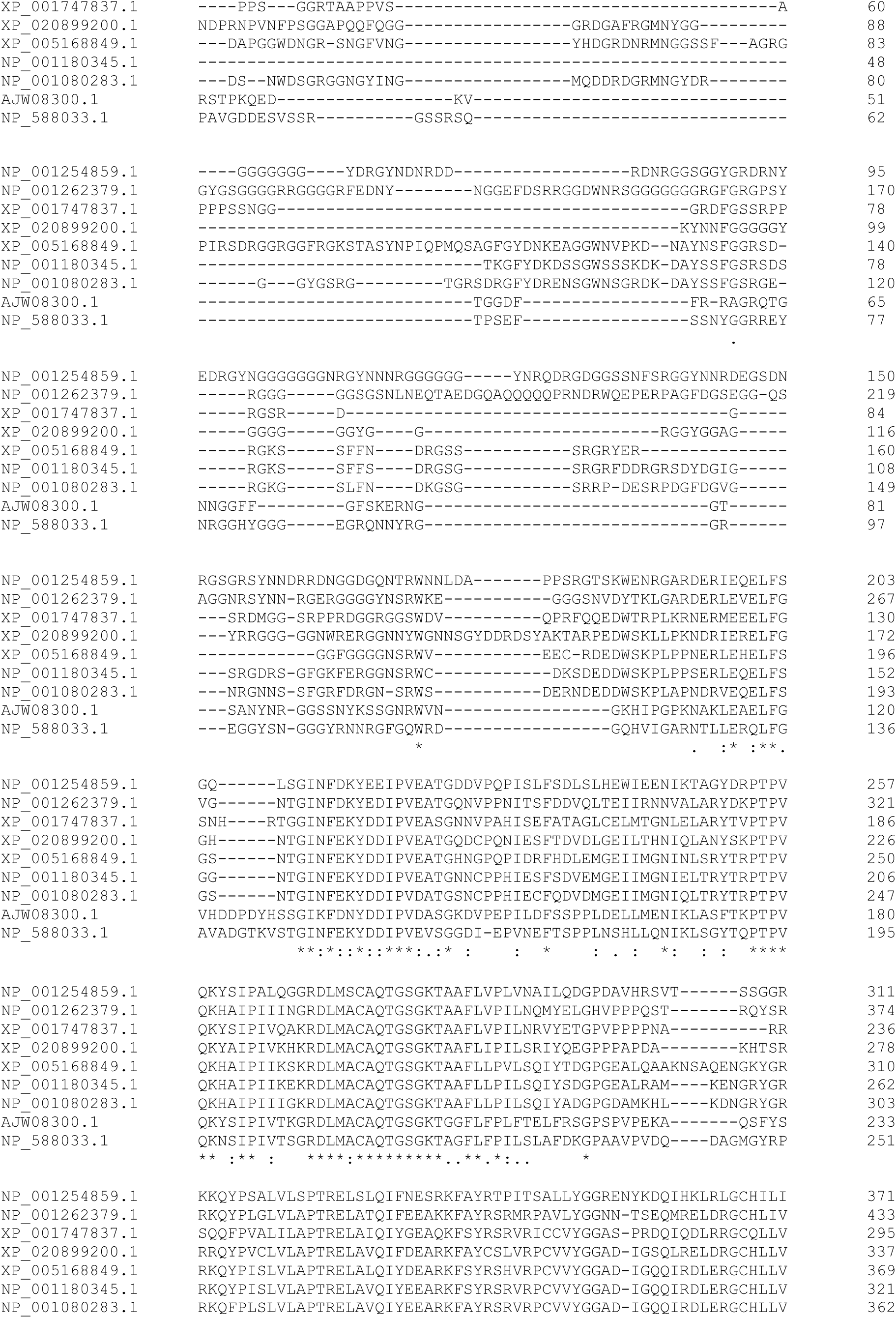
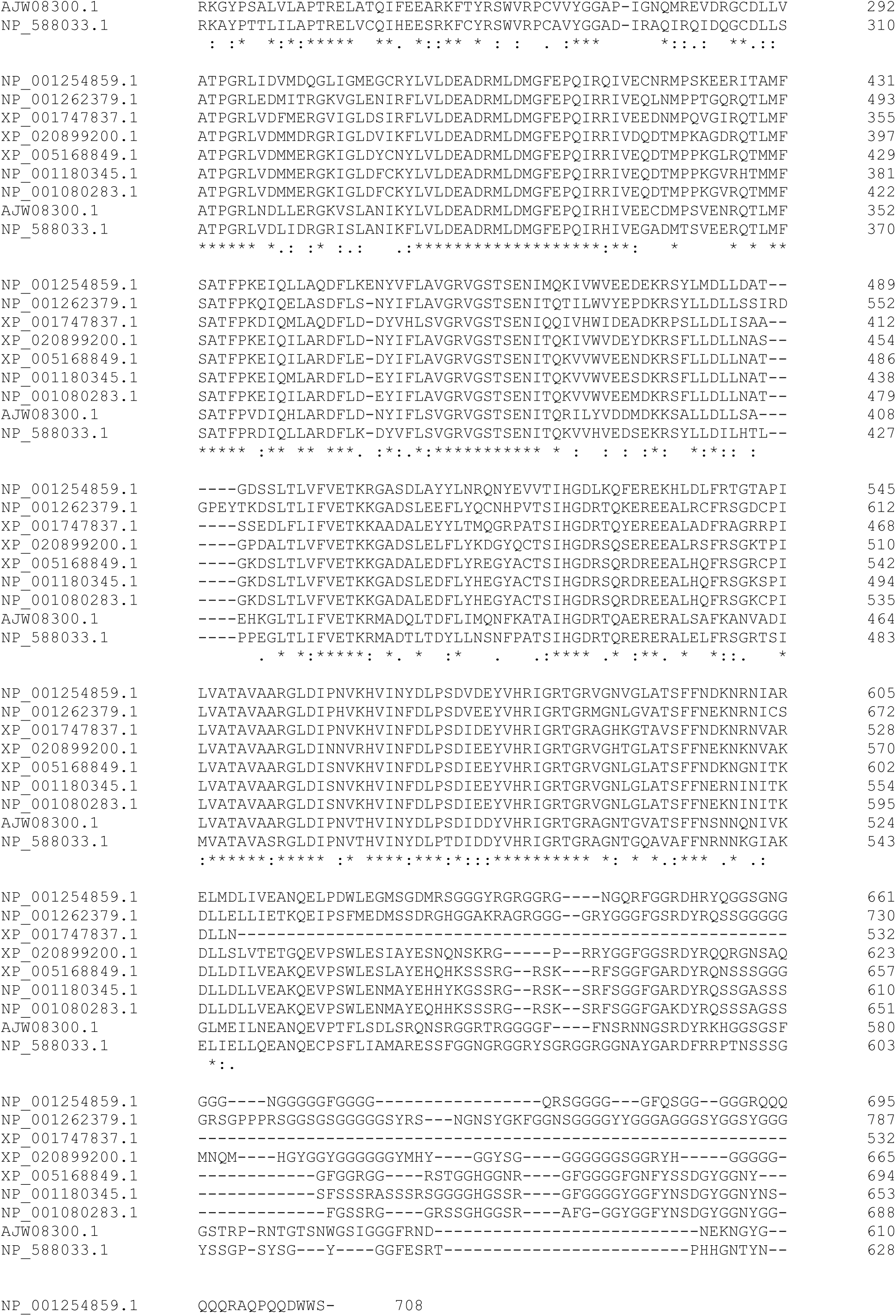
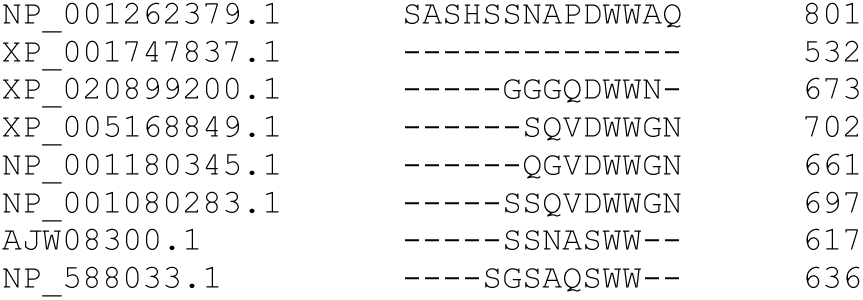

### 1.4 Calculation of minimum possible SCD for sequence with same composition as LAF-1 RGG

To obtain a sequence with the minimum possible SCD value, the charged amino acids must be clustered at the very ends of the sequence with positive charges at one end, negative charges at the other, and uncharged amino acids in between. Since we consider histidine in our model to have a +0.5 charge, the +1 charged amino acids should be at the very end with histidine residues following.

We also must consider that for in vitro studies, the initial methionine residue and the LEHHHHHH tag must be conserved. Thus a sequence with minimum possible SCD is:

MDDDDDDDDDDDDDDDDDEEE……HRRRRRRRRRRRRRRRRRRRRRRRRLEHHHHHH

With all of the uncharged residues in between the negatively-charged N-terminal, and the positively-charged C-terminal, and having an SCD of -28.032. Note that since D and E have the same charge, any permutation of residues 2-21 would not change the SCD value.

The probability of randomly sampling a sequence with the minimum SCD value can be calculated by considering the number of residues being shuffled as 176 – 9 = 167. Then one must consider the four regions that must be correct:

1. All D and E residues within 2-21
2. All R residues within 145-168
3. H residue at 144
4. All uncharged residues within 22-143

To account for these values and the degeneracies, we can calculate the probability of randomly sampling such a sequence as

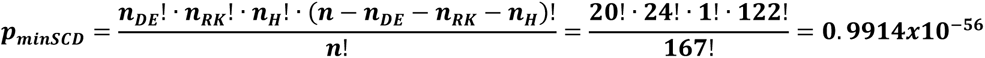

## 2. SI Figures

**Figure S1:**
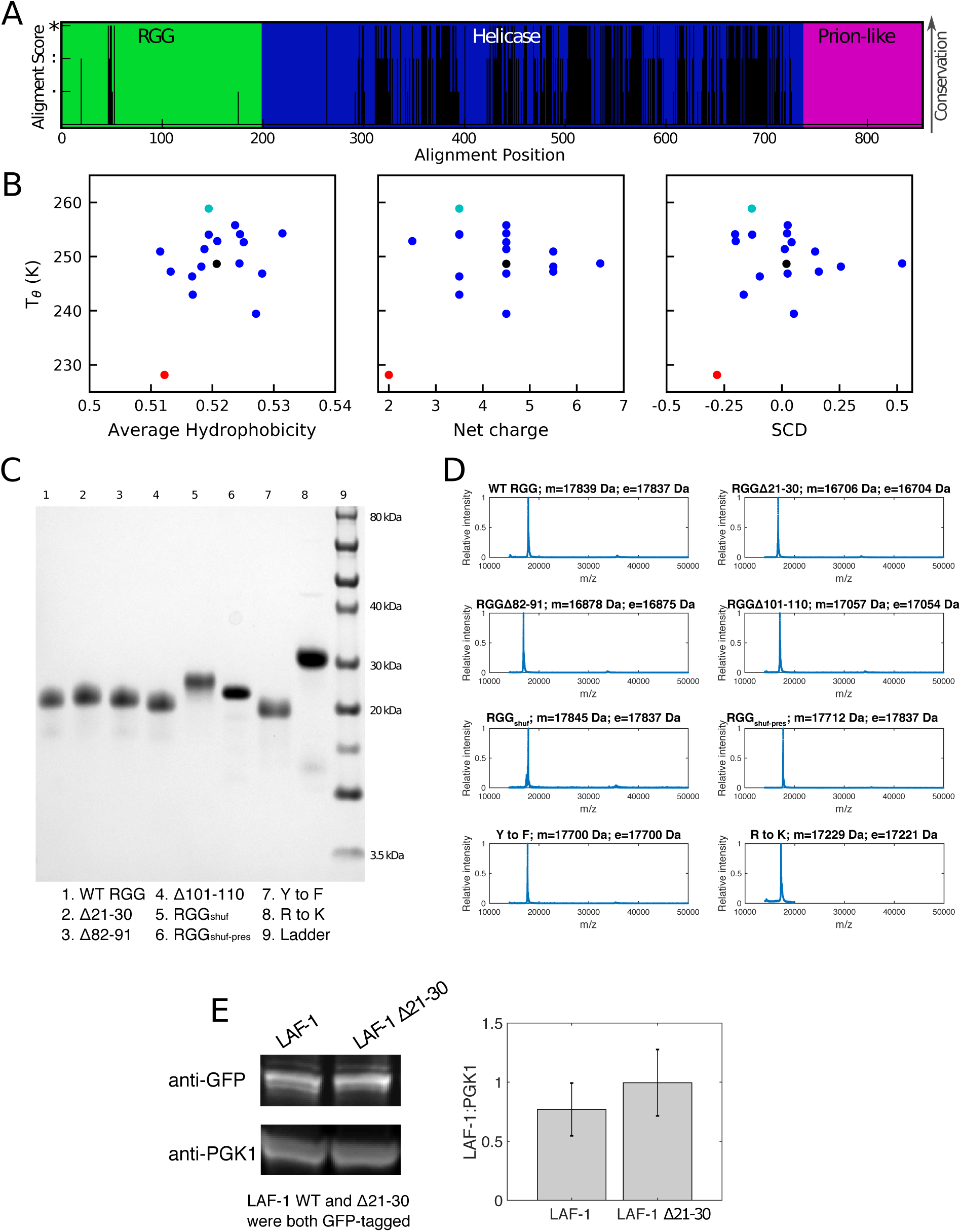
Characterization of deletion variants of LAF-1 RGG: A) Sequence conservation of full-length LAF-1, showing a high degree of conservation in folded helicase domain, and poor conservation in disordered RGG and prion-like domains. B) Tθ calculated from single-chain simulations of the deletion series (Fig. 1C) compared to sequence descriptors. In general, higher Tθ is expected to be associated with higher average hydrophobicity, smaller absolute net charge, and more negative SCD. The symbol colors correspond to WT (black), Δ21-30 (red), Δ101-110 (cyan), with all other variants represented as blue. C) SDS-PAGE gel of purified RGG and its variants. D) MALDI-TOF mass spectra of RGG domain and its variants, where m denotes measured and e denotes expected molecular mass. The only discrepancy > 10 Da is RGG_shuf-pres_ which is likely due to the loss of initiating methionine. E) Western blot shows a similar expression level of LAF-1 WT and LAF-1 Δ21-30 in yeast.

**Figure S2:**
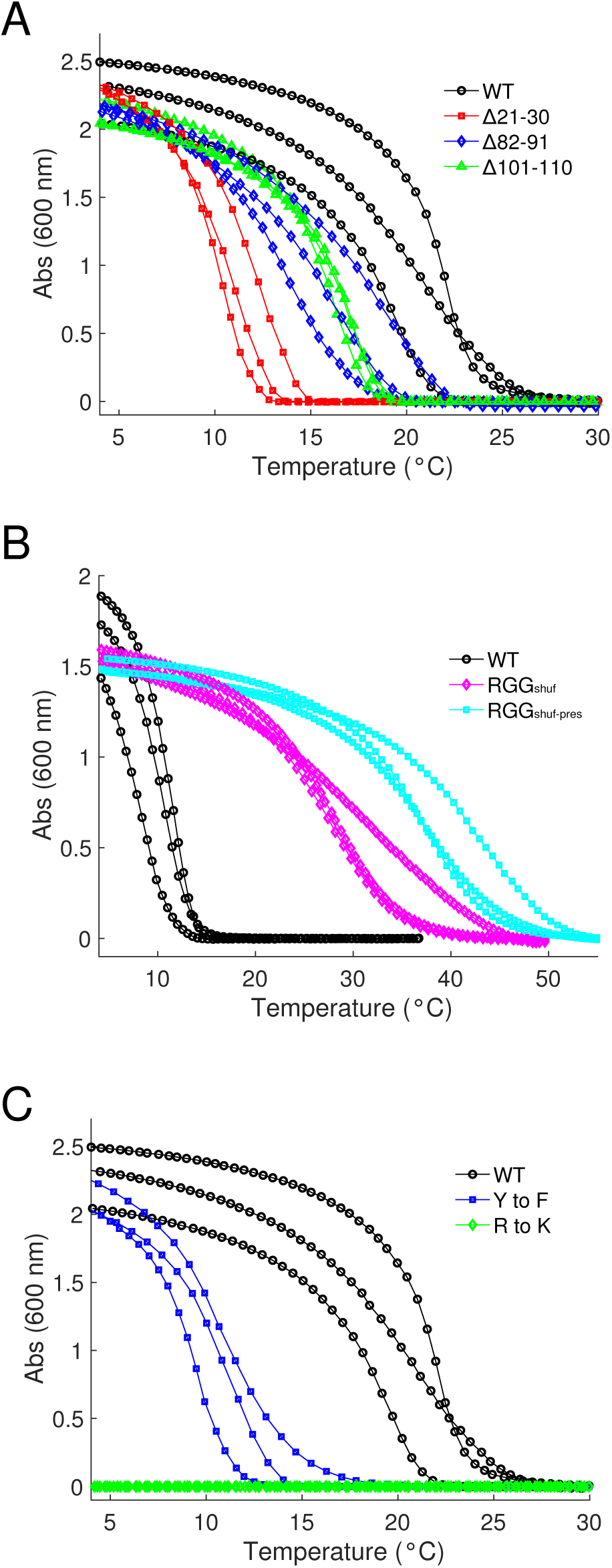
Replicates of turbidity experiments. (corresponding to Fig. 1D, 2D, 3B): Turbidity curves of WT and A) deletion variants, B) shuffled sequences, and C) bulk mutations. Protein concentrations were 1 mg/mL for (A) and (C) and 0.3 mg/mL in (B). WT data is the same for (A) and (C). In all cases, proteins were in 150 mM NaCl, 20 mM Tris, pH 7.5.

**Figure S3:**
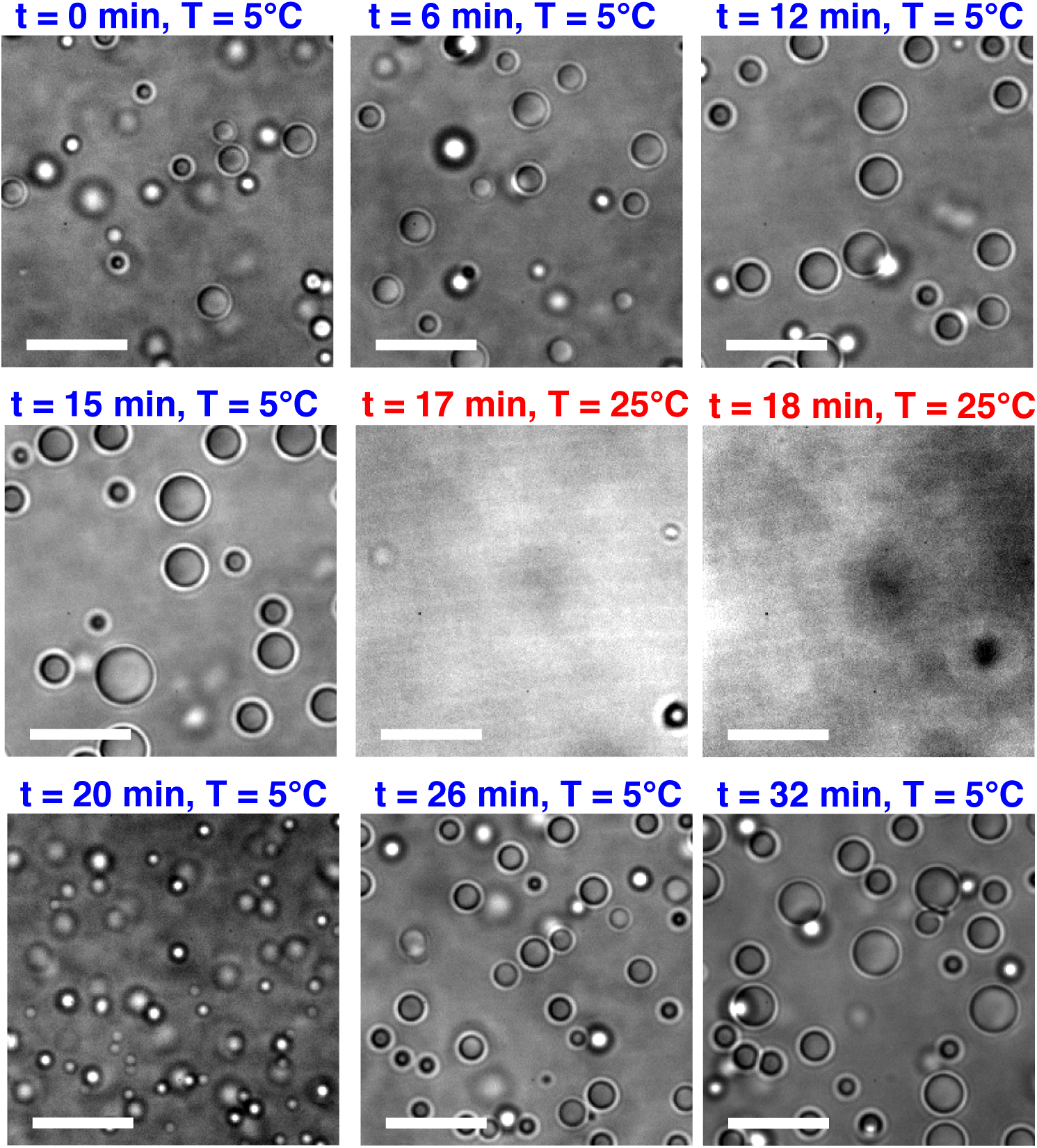
Reversible LLPS of Δ21-30 variant: The Δ21-30 variant of RGG undergoes reversible, temperature-dependent LLPS. Snapshots follow the formation of droplets over time starting at low temperature (5 °C), then rapidly increasing temperature from 5 °C to 25 °C to disperse the droplets, and then rapidly decreasing the temperature back to 5 °C to induce phase separation again. Scale bars: 10 μm.

**Figure S4:**
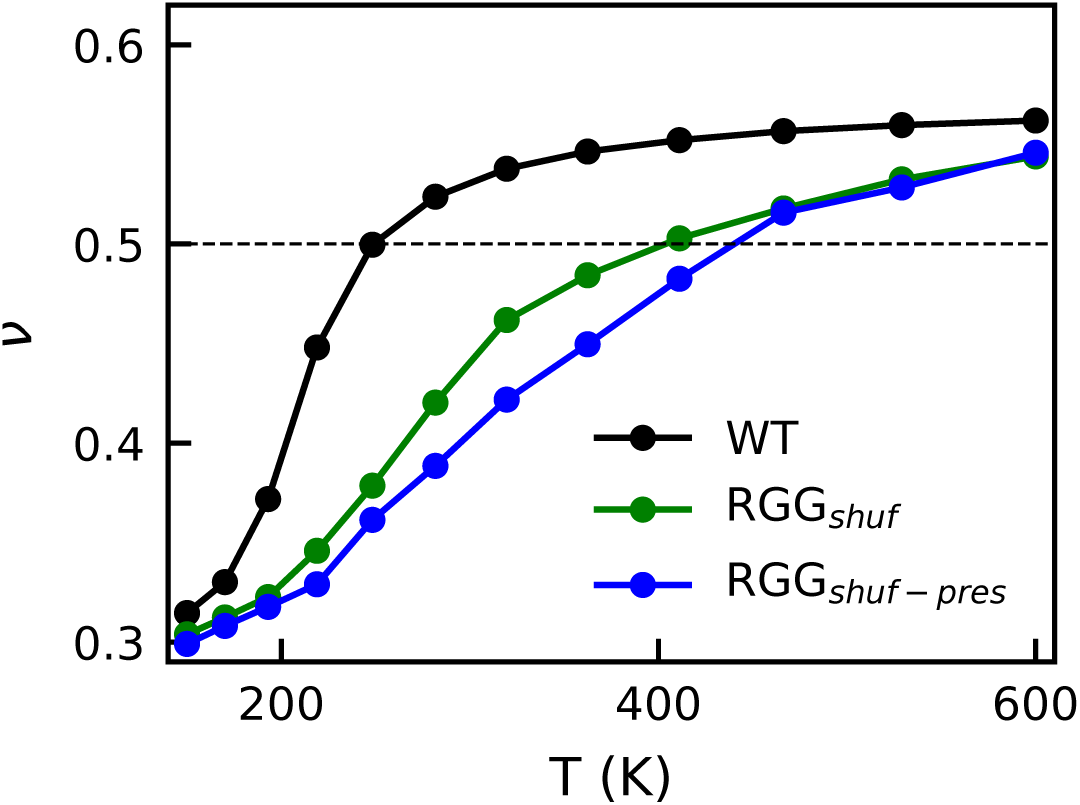
Single-chain compactness of RGG and shuffled variants: We calculate the Flory scaling exponent (ν) of the three variants of RGG as in previous work^40, 76^ and see that the WT is significantly more extended than the shuffled variants at a wide range of temperatures. We also see that RGG_shuf-pres_ is marginally more compact than RGGshuf, consistent with our experimental results showing that RGG_shuf-pres_ has the greatest LLPS propensity.

**Fig S5:**
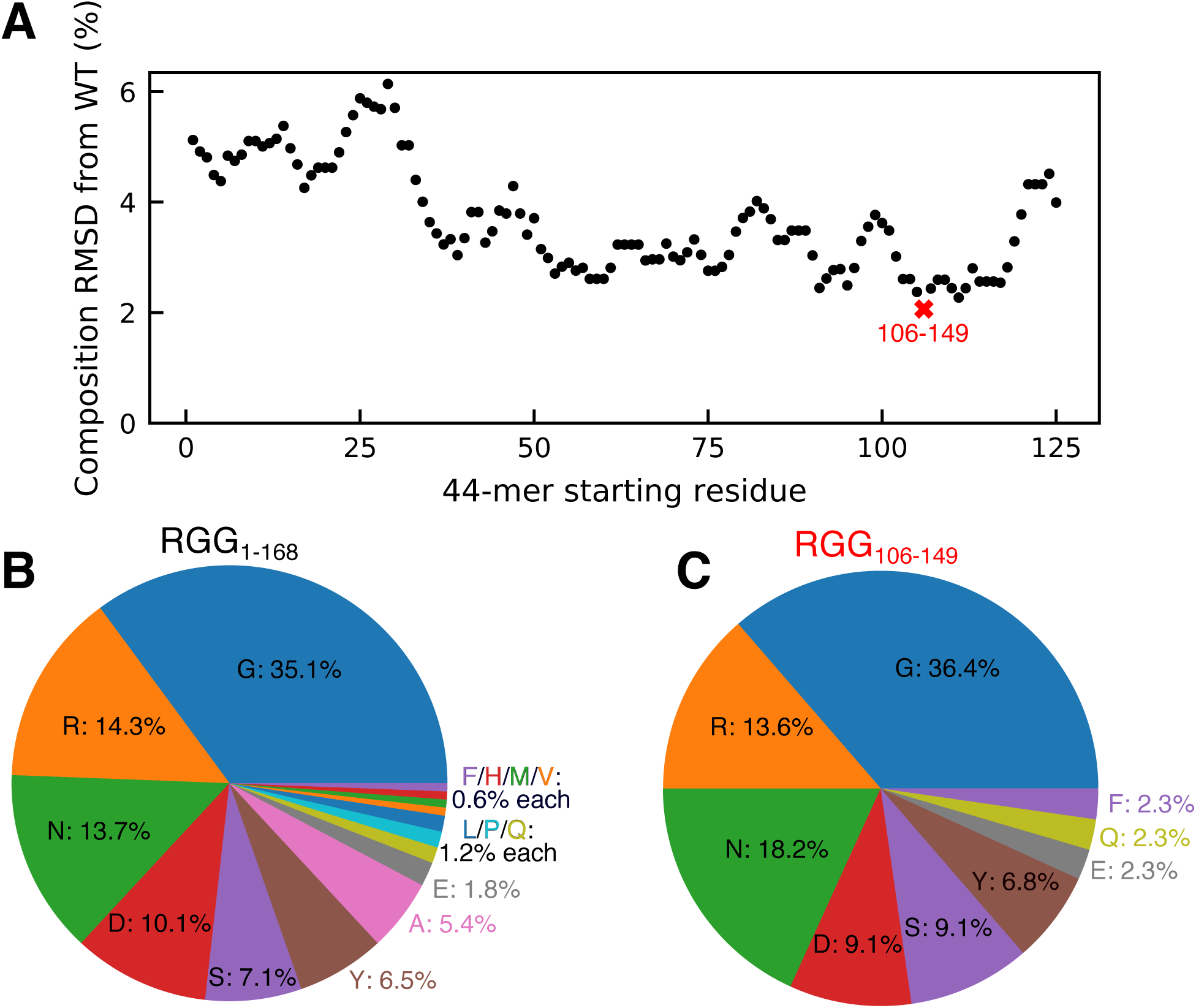
Sequence composition of WT RGG and 44-residue fragments: A) Composition-based RMSD is calculated for all continuous 44-residue fragments of LAF-1 RGG, showing the overall compositional similarity with the full 168-residue sequence. A total of 168 – 44 + 1 = 125 sequences of 44 residues were tested. B) Pie chart of amino acid composition of WT RGG is highly similar to C) pie chart of the lowest-RMSD 44-mer, RGG_106-149_.

**Figure S6:**
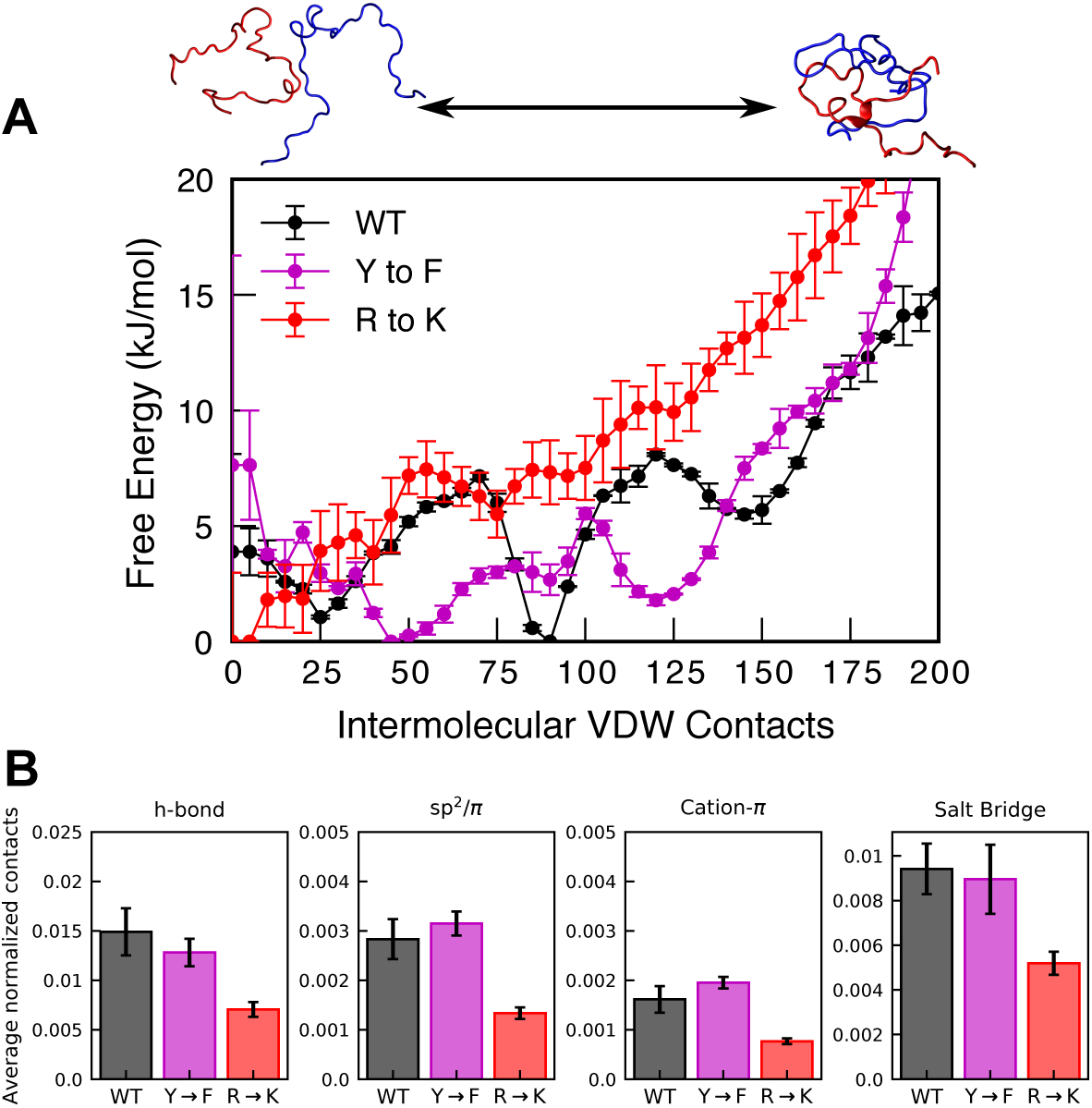
All-atom simulations of RGG106-149 show R→K has lower self-association: A) Free energy profile of contact formation between two identical RGG106-149 chains from simulations using well-tempered metadynamics. Both WT and Y→F show global minima at a finite number of contacts, while R→K has a global minimum at 0 contacts, indicating unfavorable self-interaction. B) Average total number of intermolecular contacts from two-chain simulations normalized by the average total number of VDW contacts for that system. Errorbars for all plots are SEM with n = 2.

**Figure S7:**
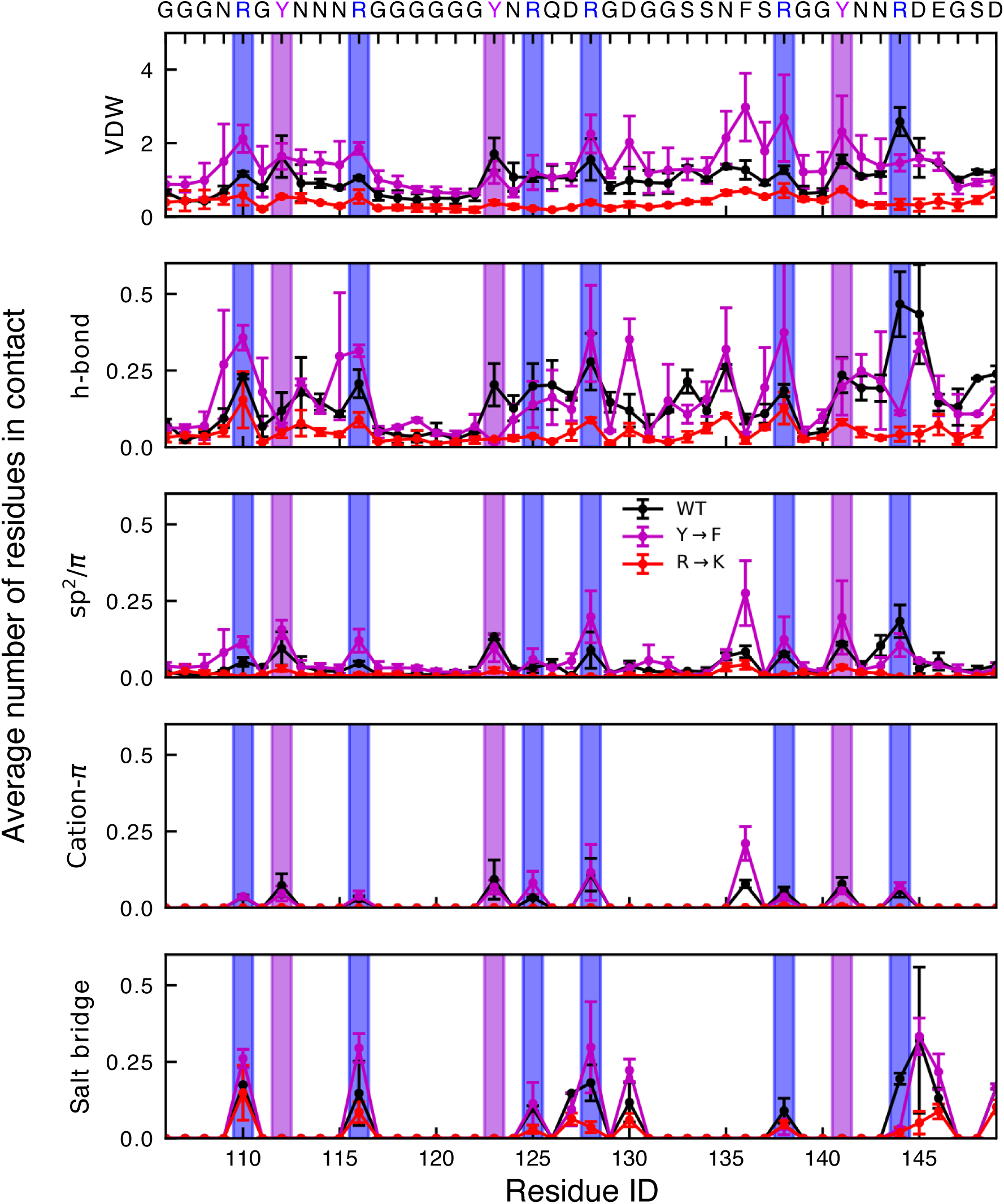
Per-residue contacts from all-atom simulations: Average number of intermolecular residue-residue pairs for each residue of the RGG106-149 sequence. Two residues are considered to be in contact if there is at least one atom from each residue in contact (VDW) or at least one hydrogen bond, sp^2^/π interaction, cation-π interaction, or salt bridge between the two residues. In the case of VDW, multiple residues may be in contact with a single residue as only one atom needs to be in contact, and residues may have VDW interactions with many other amino acids on the other protein chain. Generally, we see that VDW interactions and hydrogen bonds are well-distributed throughout the sequence for all variants of RGG106-149. Cation-pi, sp^2^/π, and salt-bridge interactions are less well-distributed due to their dependence on certain amino acid side chains. To highlight the contribution of aromatic and cationic residues, we have highlighted the arginine and tyrosine residues in these plots. Error bars are SEM with n = 2.

**Figure S8:**
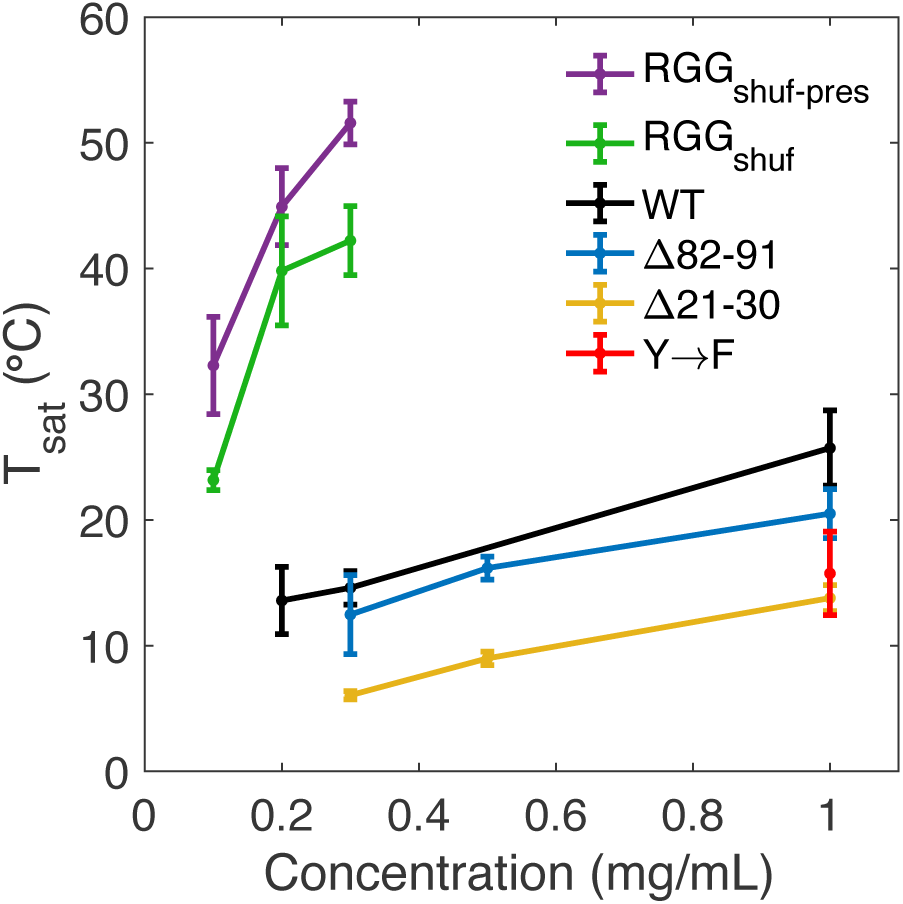
Phase diagrams of RGG mutants: Saturation temperature as a function of total protein concentration for the two shuffled variants, two deletion variants, Y→F, and WT RGG. T_sat_ values are determined from turbidity curves where absorbance first exceeds 0.02. Error bars are STD with n = 3. T_sat_ of WT is significantly different than that of Δ21-30, RGGshuf, and RGG_shuf-pres_ (p ≤ 0.005), but not significantly different than that of Δ82-91 (p = 0.73), based on one-way ANOVA followed by Tukey’s post-hoc test at 0.3 mg/mL. T_sat_ of WT is significantly different than that of Δ21-30 and Y→F (p ≤ 0.005), but not Δ82-91 (p = 0.12), based on one-way ANOVA followed by Tukey’s post-hoc test at 1 mg/mL.

**Figure S9:**
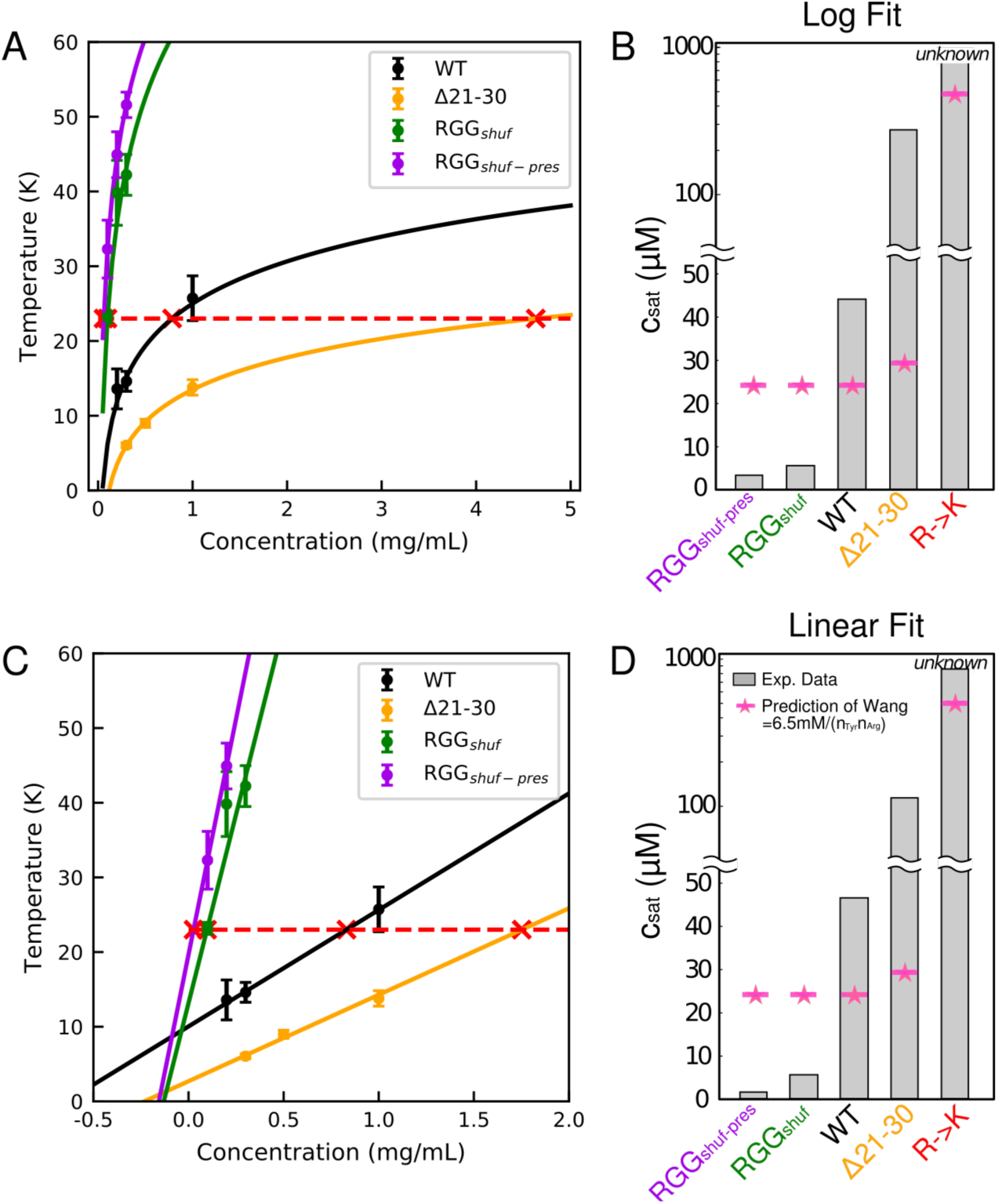
Fitting of c_sat_ from experimental data: A) Logarithmic fits to experimental data to calculate c_sat_ (red X’s) at 23°C, (red dashed line). B) bar plot of saturation concentrations for the different variants of RGG and comparison to empirical predictions using relationship from Wang et al. C) Linear fits to experimental data to calculate c_sat_ as before. For RGG_shuf-pres_, one data point was removed from the fitting such that the extrapolated c_sat_ value would be positive. D) bar plot of saturation concentrations for the different variants of RGG using the linear fit and compared to empirical predictions using relationship from Wang et al. Error bars are STD with n = 3.

**Figure S10:**
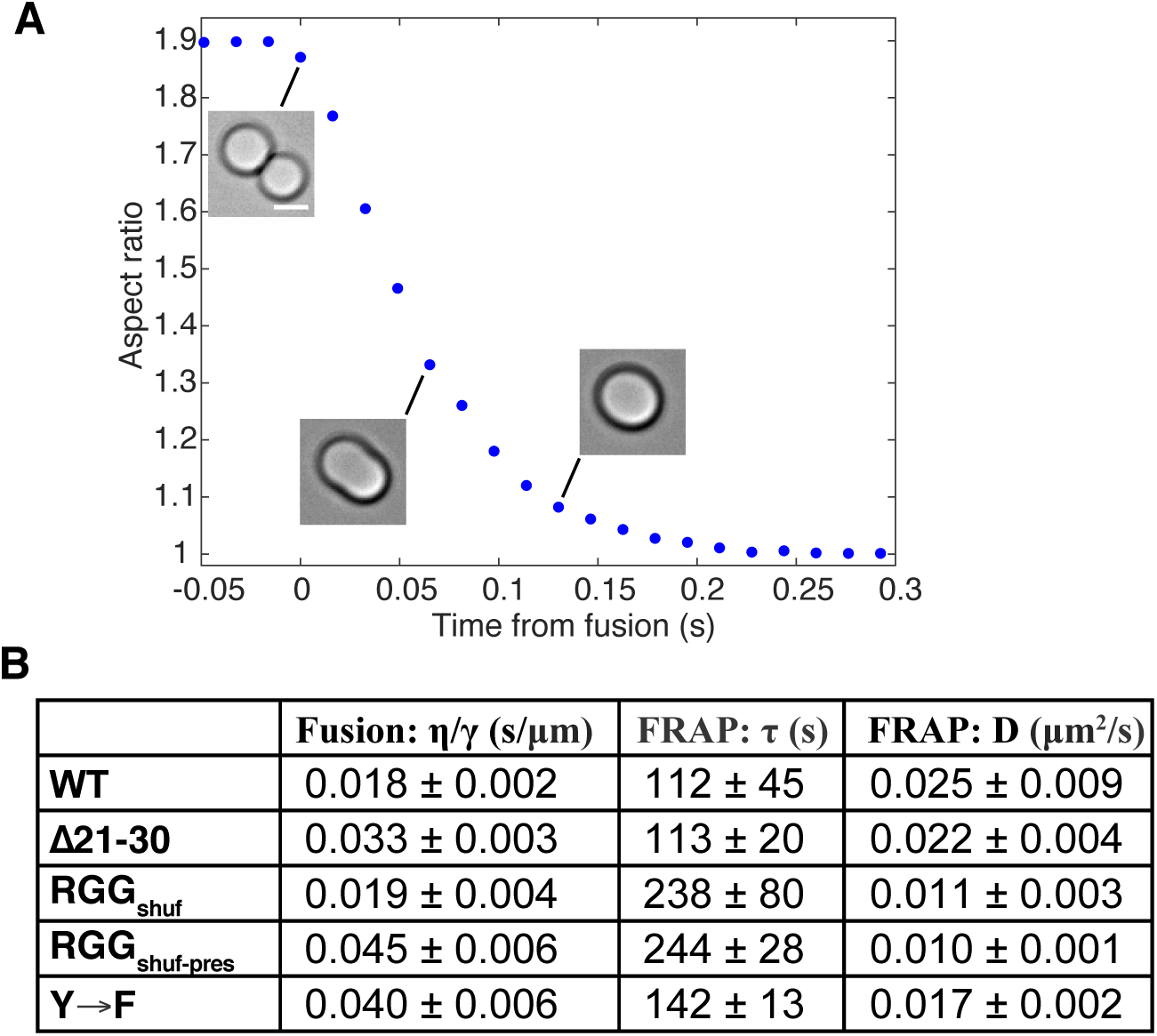
Measurements of droplet material properties: A) Example trace showing aspect ratio of fusing droplets relaxing exponentially to a sphere, from which the relaxation timescale is calculated. The data shown corresponds to the Y→F droplet fusion event in Fig. 5A, several images of which are reproduced here as insets beside their corresponding data points (scale bar: 2 μm). B) Table summarizing measurements of inverse capillary velocity η/γ from droplet fusion experiments, as well as recovery timescale τ and diffusivity D from FRAP.

